# Structure of the Hibernating *Francisella tularensis* Ribosome and Mechanistic Insights into Its Inhibition by Antibiotics

**DOI:** 10.1101/2025.11.25.690415

**Authors:** Martin Klima, Jan Silhan, Pavla Pavlik, Kamil Hercik, Evzen Boura

## Abstract

*Francisella tularensis* is the causative agent of tularemia, a zoonotic disease named after the city of Tulare, California. Symptoms include sudden fever, chills, fatigue, and swollen lymph nodes, among others, and without treatment it is very serious or even fatal. In addition, *F. tularensis* is considered a potential bioterrorism threat due to its high infectivity and lethality. Ribosomes are key targets for many classes of antibiotics. In this study, we examined the *F. tularensis* ribosome and determined its structure at 2.8Å resolution using cryo-electron microscopy. Notably, we observed the stress-induced ribosome-associated inhibitor A (RaiA) protein bound to the ribosome. RaiA functions as a molecular hibernation factor, inhibiting bacterial translation in response to stress or nutrient deprivation. This mechanism parallels that described in the model organism *Escherichia coli* and in several pathogenic bacteria, such as *Staphylococcus aureus.* Furthermore, we solved structures of the antibiotics chloramphenicol and gentamicin bound to the *F. tularensis* ribosome. Collectively, these results provide structural insights that highlight previously unexplored opportunities for therapeutic intervention.

## Introduction

*Francisella tularensis* is a Gram-negative, highly infectious, rod-shaped intracellular bacterium first isolated by McCoy and Chapin (1912) from rodents in Tulare County, California. It is the causative agent of tularemia, a fulminant and debilitating zoonotic disease in humans. Low infectious dose (∼25 colony-forming units), ease of dissemination and aerosolization, potential high mortality rate, and the need for a public health response [1] led CDC to categorize the agent in Category A, the highest-priority group of biological agents and toxins in relation to the risk of bioterrorism. To date, four subspecies of *F. tularensis* have been identified, exhibiting distinct biochemical properties, virulence features, and geographic distributions [2]. The highly virulent *F. tularensis* subsp. *tularensis* (type A), found only in North America, has been further divided into subtypes A.I and A.II based on differences in virulence [3]. The moderately virulent *F. tularensis* subsp. *holarctica* (type B) is endemic in the Northern Hemisphere [4]. Strains of *F. tularensis* subsp. *mediasiatica* have been isolated only in a region of Central Asia [2]. The least virulent, opportunistic *F. tularensis* subsp. *novicida* appears to be distributed throughout the United States, with some reported isolations from Australia [5]. Due to the high threat to public health systems, the absence of a licensed vaccine, and the potential for complicated treatment, the significant risk of deliberate misuse as a bioterrorism agent with numerous casualties and widespread panic—leading to devastating effects on the economy and public safety—makes the search for new drug targets and treatment approaches highly urgent.

Ribosomes are central and complex molecular machines for protein synthesis and, as such, are critical components of any cell. Not surprisingly, bacterial ribosomes are targets of many important antibiotics used in human and veterinary medicine. Among the drugs currently used to treat tularemia are several compounds that target the ribosome [6]. Streptomycin, gentamicin, and tetracycline bind the 30S subunit, whereas chloramphenicol binds the 50S subunit. Streptomycin and gentamicin are preferred for severe tularemia disease and are recommended by CDC [7]. Of course, antibiotics targeting other biochemical pathways, such as fluoroquinolones (e.g., ciprofloxacin), may also be used [8, 9]. Nevertheless, the ribosome is a prime target for pharmacological intervention not only in the case of *F. tularensis*.

Protein synthesis, carried out by ribosomes, is the most energy-consuming process in the cell [10]. As a survival strategy, organisms have evolved strategies to switch ribosomes off when not needed (e.g. stationary phase) or under stress (starvation or low temperature). This is facilitated by ribosome-sleeping, or hibernation, factors. Notably, several families of these factors exist and their molecular mechanisms differ, yet the outcome is a “sleeping” ribosome. For example, HPF (hibernation-promoting factor) promotes dimerization of ribosomes into a catalytically inactive 100S dimer [11, 12]. The recently described Balon factor binds the ribosomal A site in an mRNA-independent manner and promotes hibernation of the ribosome [13]. Similarly, RaiA binds near the A site to block tRNA entry efficiently switching off translation [14]. In addition, the hibernation factors protect "sleeping" ribosomes from degradation by endonucleases [15, 16].

In this study, we describe the structure of the *F. tularensis* ribosome in a hibernating state with RaiA bound, as well as its complexes with the antibiotics chloramphenicol and gentamicin. These structures reveal conserved sites relevant for drug design and suggest that the RaiA binding site may represent a druggable pocket.

## Results

### Ribosome preparation

Several *F. tularensis* subsp. *tularensis* strains (including the Schu S4 strain) are highly pathogenic and are classified as biosafety level 3 (BSL-3) agents, depending on national regulations [17]. We aimed to structurally characterize its ribosome. To achieve this, we took advantage of the fact that the well-characterized *F. tularensis* subsp. *holarctica* strain FSC200, isolated in Sweden [18] is classified as BSL-2 in the European Union [19]. Notably, its ribosome shows an almost perfect match to that of the highly virulent Schu S4 strain (SI Figure 1). This allowed us to cultivate the FSC200 strain safely under BSL-2 conditions and to obtain sufficient material for ribosome purification. The purification procedure itself is described in detail in the Materials and Methods section.

**Figure 1.**
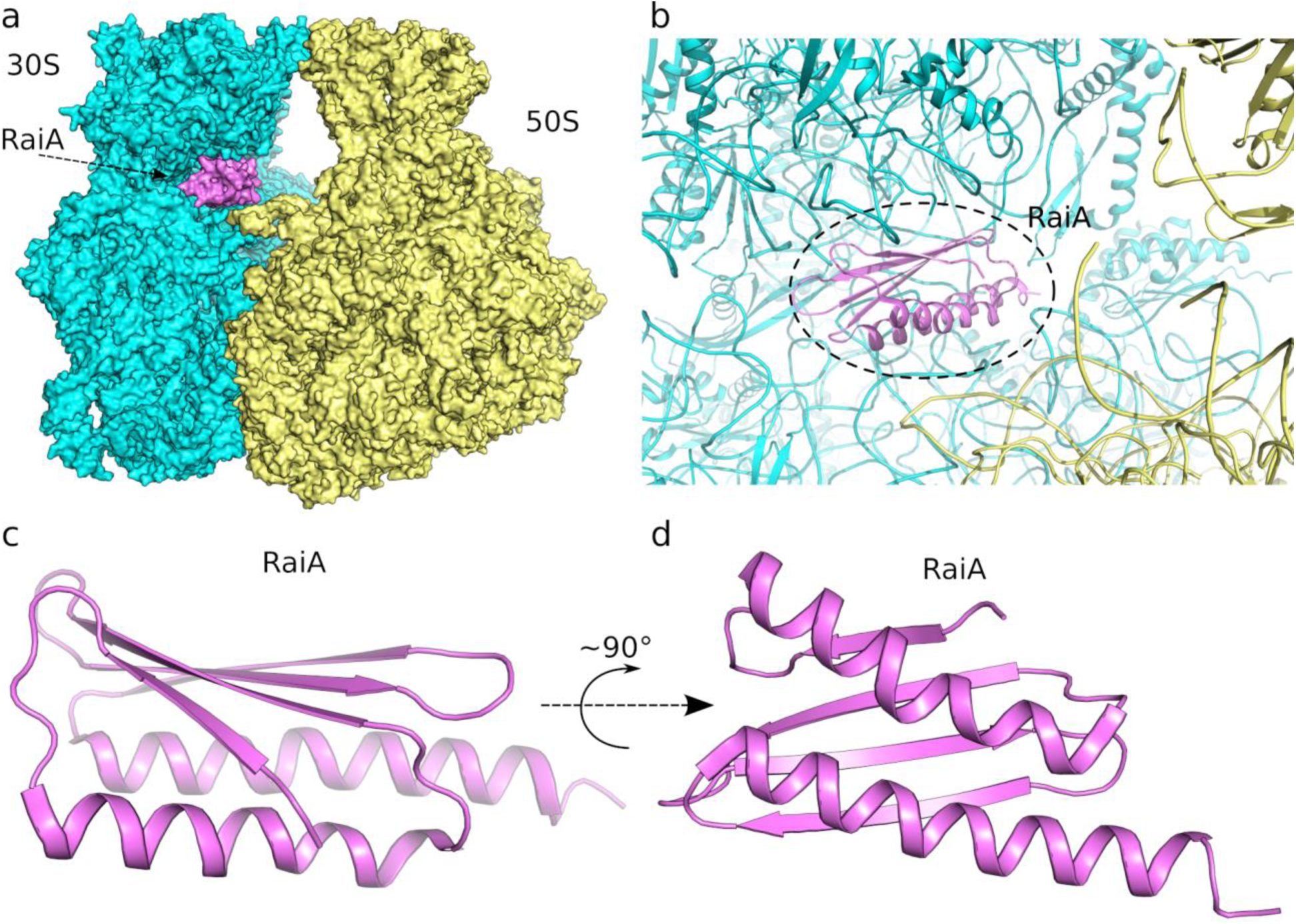
Overall architecture of the *F. tularensis* 70S ribosome and the location of RaiA. Surface representation of the *F. tularensis* 70S ribosome showing the small 30S subunit (cyan), large 50S subunit (yellow), and the ribosome-associated inhibitor A (RaiA, violet). (b) View of RaiA bound within its specific site on the 70S ribosome at the intersubunit interface. (c, d) Detailed cartoon representations of RaiA highlighting its overall fold and secondary structure elements.

### Structure of F. tularensis ribosome

To obtain the structure of the *F. tularensis* ribosome, we used the cryo-EM method. We collected 28,005 movies using the Titan Krios microscope which allowed us to refine the cryo-EM map at 2.5 Å using 476,383 particles to solve the structure (SI Figure 2). To build the initial model, we used a 70S ribosome structure of *Staphylococcus aureus* (pdb entry 5LI0) [20] as a template. The sequences of the 23S, 16S, and 5S ribosomal RNA molecules of *S. aureus* were aligned to those of *F. tularensis,* and their sequences were changed accordingly. Next, we used AlphaFold3 models for the proteins that we expected to be present in the 70S ribosome (29 large-subunit and 19 small-subunit proteins), aligned them with the template model, placed them within the experimental density maps, and further refined them. Finally, proteins absent from the *S. aureus* template but present in *F. tularensis* ribosomes were fitted manually, as described below.

**Figure 2.**
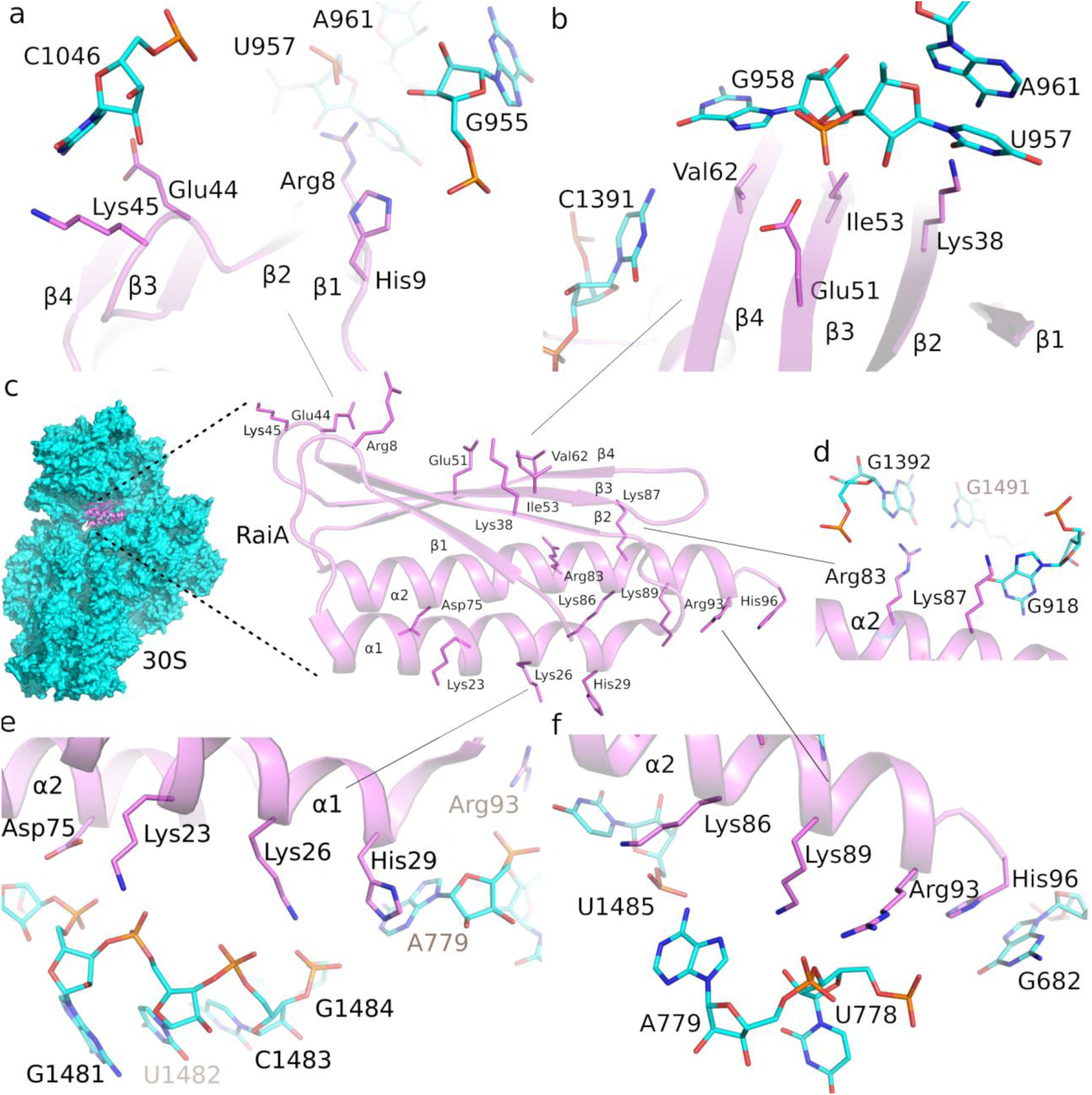
Details of molecular interactions between RaiA and the 30S subunit of *F. tularensis*. (a, b, d–f) Close-up views of residues mediating contacts between RaiA (violet) and the 16S rRNA of the 30S subunit (cyan). For clarity, only residues within interaction distance are shown as sticks. Carbon and heteroatoms are coloured according to their respective chains. (e, f) Two rows of highly conserved charged residues located on helices α1 and α2 interact predominantly with the phosphate backbone of the 16S rRNA.

Very soon in the refinement process we noticed an excess electron density that did not correspond to any of the proteins we expected in the *F. tularensis* 70S ribosome. This unknown density was initially modelled with a poly-alanine chain, and the presence of distinct side-chain densities enabled a reasonably precise fit of 21 bulky amino acids. The resulting structure was analyzed using the Foldseek server [21] which returned an unambiguous match to the ribosome hibernation-promoting factor RaiA. Its AlphaFold model was placed in the experimental map and further refined as above (SI Figure 3).

**Figure 3.**
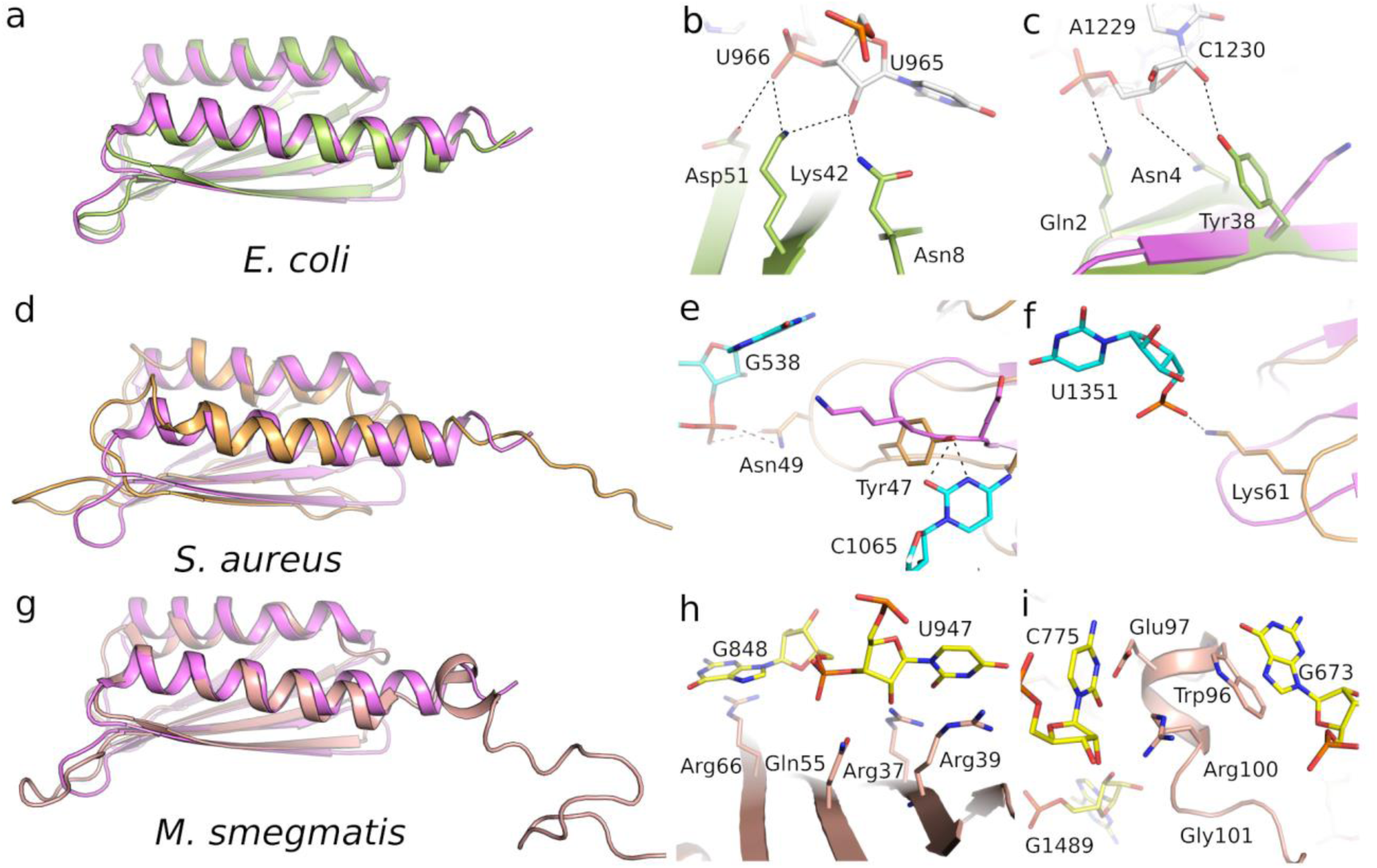
Comparative structural features of the N-terminal region and RNA-binding sites across species. (a, d, g) Superimposed structures of the *E. coli* (green), *S. aureus* (orange), *Mycobacterium* (pink), and *F. tularensis* (violet) RaiA orthologues, highlighting overall fold conservation and species-specific differences. (b, c) *E. coli* (PDB ID: 6H4N), (e, f) *S. aureus* (PDB ID: 6S0X), and (h, i) *Mycobacterium smegmatis* (PDB ID: 8WHX) structures showing detailed views of prominent interaction differences among ribosome hibernation factors and their orthologues from the selected organisms.

RaiA occupies a site at the intersubunit interface adjacent to the mRNA channel and the decoding centre. The electron density was very well defined and allowed us to not only to build *F. tularensis* RaiA but also to place side chains confidently, especially at the RaiA–rRNA interface, revealing the detailed network of contacts with helices and loops of the 16S rRNA that stabilises the inactive 70S monomer. (**Figure 1**).

### RaiA: structure and binding to the ribosome

*F. tularensis* RaiA is composed of four β-sheets and two α-helices arranged in the order β1–α1–β2–β3–β4–α2, which is consistent with previously solved structures of RaiA from related bacteria [22–27]. The protein interacts exclusively with the 16S rRNA of the 30S subunit. Charged residues Arg8 and His9, located in the β1–α1 loop, lie near the A-site tRNA binding region, where Arg8 interacts with U957 and His9 contacts the phosphate of G955. In the β3–β4 loop, Glu44 and Lys45 are positioned to interact with nucleotide C1046 (**Figure 2a**).

A cluster of residues lying along the β-sheet plane—Val62, Glu51, Ile53, and Lys38—engages rRNA bases C1391 and the patch formed by G958, U957, and A961. Helix α1 contains basic residues Lys23, Lys26, and His29, which, together with Asp75 from helix α2, interact with the rRNA backbone between A1480 and G1484 (**Figure 2b**).

Helix α2, positioned at the P-site of the 30S subunit, contains two positively charged residues, Arg83 and Lys87, that contact G1392 and G918, respectively. Oriented in the opposite direction from the P-site, another group of positively charged residues—Lys86, Lys59, and Arg93—interacts predominantly with the phosphodiester backbone of U1485, A779, and U778 (**Figure 2c-f**). At the C-terminus, His96 forms a π–π interaction with G692 (**Figure 2f**), a contact commonly conserved among RaiA homologues [24, 26]. Detailed cryo-EM maps documenting these interactions (**Figure 2**) are shown in **SI Figure 4**.

**Figure 4.**
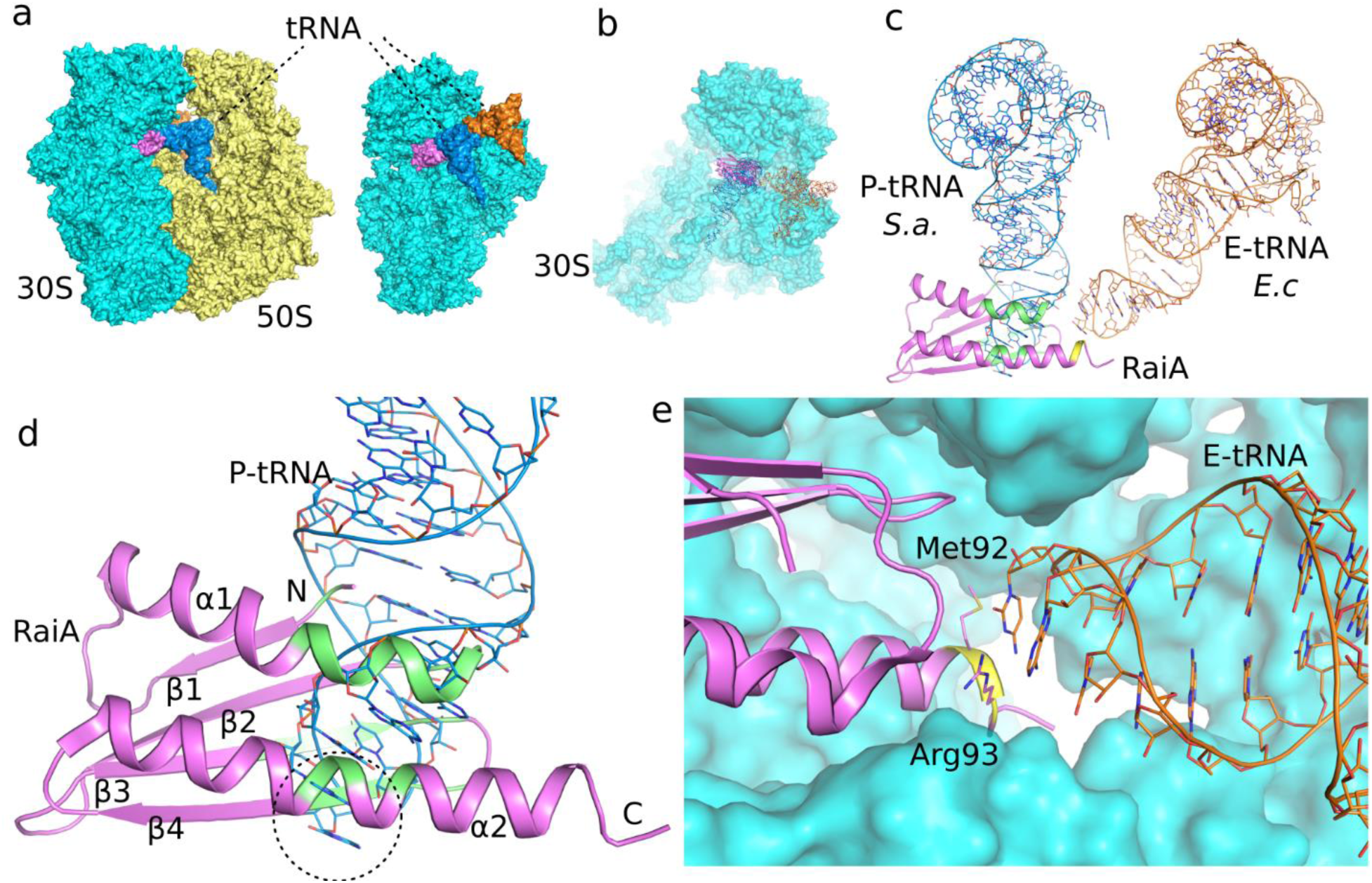
Structural basis of RaiA positioning on the *F. tularensis* ribosome and its overlap with tRNA-binding sites. (a) Cryo-EM structure of the *F. tularensis* 70S ribosome showing the 30S subunit (cyan), 50S subunit (yellow), and RaiA (violet), with P-site and E-site tRNAs modelled based on PDB ID: 7NMH and PDB ID: 6H4N, respectively. (a) The isolated 30S subunit highlighting RaiA (magenta) positioned in the intersubunit space, overlapping with canonical P-site (blue) and E-site (orange) tRNA positions derived from previously solved ribosome structures. (b) Overlay of the *F. tularensis* 70S–RaiA complex with P- and E-site tRNAs showing steric overlap (highlighted in green for the P-site and yellow for the E-site), indicating that RaiA occupies a position that would clash with both tRNAs. (d) Close-up view of the RaiA–P-site tRNA interface. The β-sheet and α-helical elements of RaiA (green) extend into the P-site tRNA anticodon loop (dotted circle). (e) Detail of the modelled RaiA–E-site tRNA interface within the 30S subunit (cyan surface). The region of RaiA (magenta) is in close proximity to, but not clashing, with the E-site tRNA (orange), as the interactions are limited to two amino acid residues, Met92 and Arg93, within the interaction distance (yellow).

### Structural comparison of RNA-contact regions across species

In *F. tularensis*, Arg8 forms a hydrogen bond with the phosphodiester backbone between U957 and A956, while His9 is likely to provide an additional hydrogen bond to the backbone of G955 or A956. The equivalent Arg residue is present only in *Mycobacterium* genus, where it instead engages in a cation–π stacking interaction with U947 (**SI Figure 5a**). In *S. aureus*, Arg8 and His9 are replaced by Asp9 and Asn10, which interact with the 16S rRNA (**SI Figure 5b**) [22]. No clear contacts are observed in *M. smegmatis*, consistent with the lack of side-chain residues in interaction distance of 16S [26]. His29 in *F. tularensis* interacts with the N3 atom of A779, and its backbone carbonyl likely engages the N9 position of the same base. Despite the poor conservation of the amino acid identity at this position, the backbone–RNA contact appears to be maintained in *E. coli* [24]. In contrast, the shorter loop of *S. aureus*, which contains a Tyr residue, lacks either of these interactions, indicating that the contact is specific to longer-loop variants. Interestingly, while Lys38 in *F. tularensis* forms hydrogen bonds with both O2 of U597 and N6 of A961. In *E. coli*, the corresponding Tyr38 instead hydrogen bonds with the O2′ atom of C1230, located in a distinct 16S rRNA loop. The repositioning of this interaction site in *S. aureus* and *Mycobacterium* suggests a divergence in the local RNA-binding geometry despite conservation of the loop backbone. In *F. tularensis*, Ile52 and Val62 form a hydrophobic surface facing the base plane of G958, establishing a stabilizing nonpolar interface. Other species display shorter or reoriented side chains in this region, resulting in the replacement of the hydrophobic patch by weaker, less extensive contacts (**Figure 3**).

**Figure 5.**
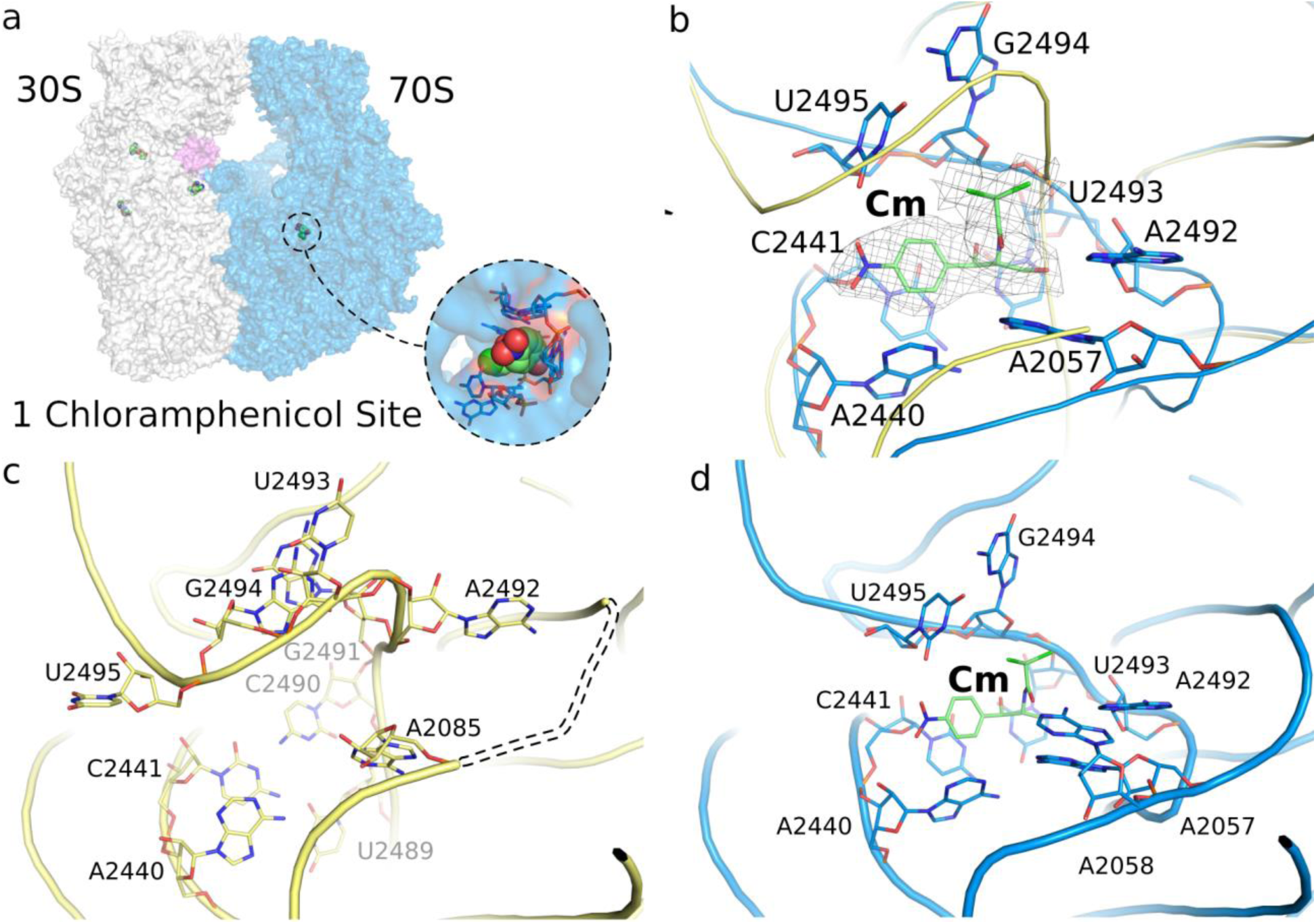
Chloramphenicol induced conformational changes in the peptidyl transferase center (PTC) (a) Surface outline of 70S ribosome (30S white and 50S blue, RaiA violet) with chloramphenicol and gentamicins bound. The chloramphenicol binding site is highlighted. (b) Overlay of the 23S rRNA of the 50S ribosomal subunit (yellow without and blue with chloramphenicol), preventing peptide bond formation. Cryo-EM density of chloramphenicol is shown as mesh. (c) Residues of the PTC before chloramphenicol binding, unmodeled residues due to poor density in the region are dashed. (d) Detailed view of 23S RNA residues responsible for chloramphenicol binding. Loop U2489–G2491 is completely displaced upon chloramphenicol binding. The previously unseen, most likely flexible A2057 (panel c), is clearly employed in the interaction with this antibiotic.

### The RaiA core is conserved, but its interaction with the 30S subunit is not

Structural alignment of *F. tularensis* RaiA with homologues from representative bacteria (*E. coli* PDB ID: 6H4N, *S. aureus* PDB ID: 6S0X, *B. subtilis* PDB ID: 5NJT, *M. smegmatis* PDB ID: 8WHX*, L innocua* PDB ID: 8UU4, and *T. thermophilus* PDB ID: 6GZQ) reveals a conserved core fold that mediates decoding-centre association (**Figure 3, SI Figure 6&7**). We show that RaiA exhibits substantial sequence and structural variation at numerous contact residues within the 30S interface, and these residues vary considerably among the species compared. Nonetheless, the high degree of conservation within the core residues is maintained across all examined examples. Besides the differences in contact residues, the *F. tularensis* RaiA orthologue lacks the extended C-terminal tail found in some species, where it modulates interactions with neighboring ribosomes or with the RMF/HPF factors that promote formation of the dormant 100S dimer [23–25]. The absence of such extensions in the *F. tularensis* RaiA correlates with the observation that *F. tularensis* ribosomes in our dataset exist largely as monomeric 70S·RaiA complexes rather than as 100S dimers, supporting a mechanistic separation between RaiA-mediated 70S stabilization and RMF/HPF-driven dimerization [23–25].

**Figure 6.**
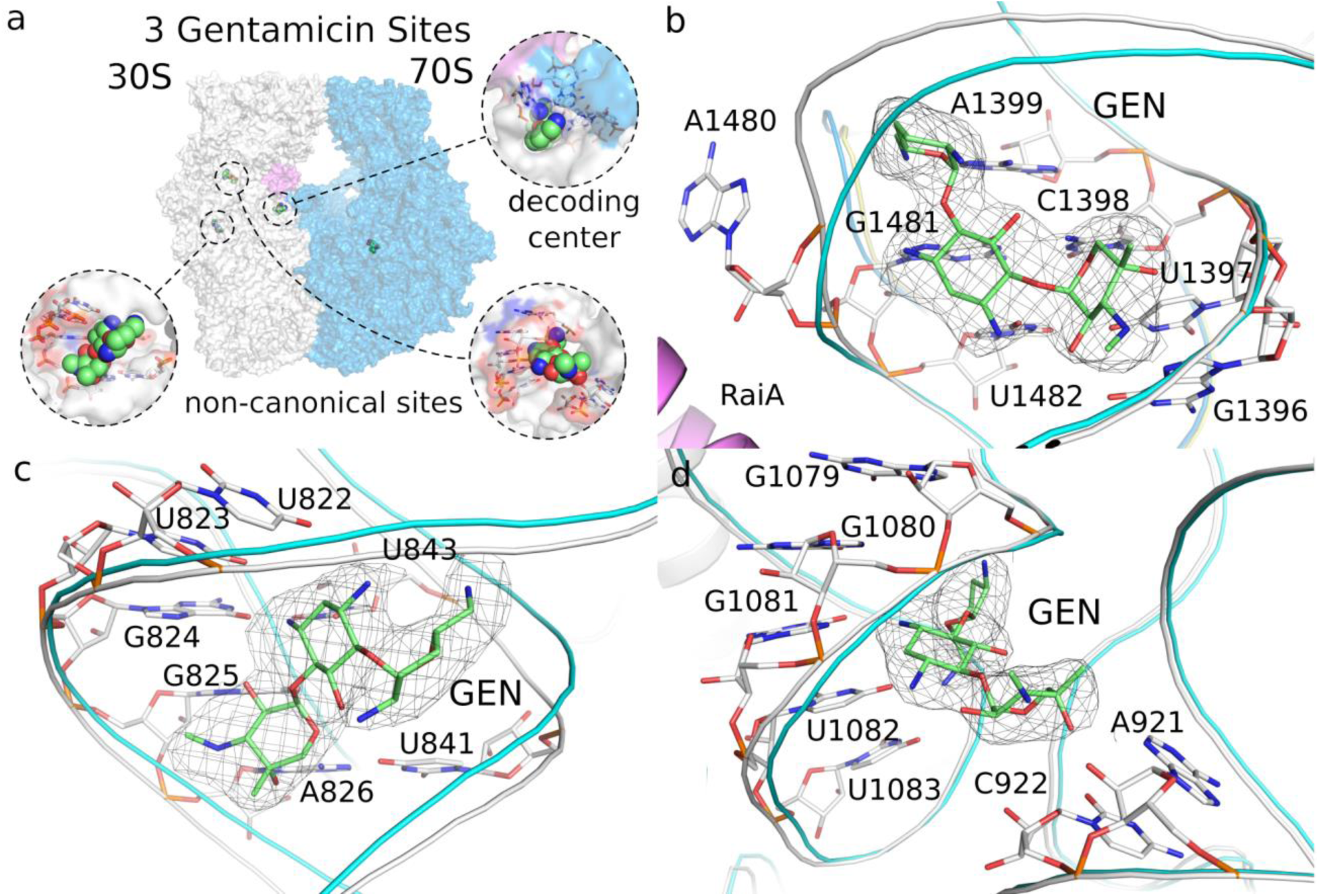
Gentamicin binds the 30S subunit of *F. tularensis* at multiple sites. (a) Surface representation of the 30S and 50S subunits showing the positions of all bound antibiotics. The three gentamicin molecules (GEN) are highlighted as spheres, with each site enlarged in the corresponding panels. (b) GEN (green sticks) bound in the canonical decoding-site pocket near RaiA and at the interface between the 30S and 50S subunits. (c) GEN bound at an additional site within the 30S subunit, interacting with residues U822-A826 with U841 and U843. (d) GEN bound at a second non-canonical site in the 30S subunit, positioned positioned simirilaly within array of five bases G1079-U1083 and two oposite bases C922 and A921. Nucleobases within 3.5 Å of each antibiotic are shown as sticks.

### F. tularensis ribosome bound to antibiotics

We also aimed to understand how the *F. tularensis* ribosome is inhibited by antibiotics. From the common antibiotics that are known to be effective against *F. tularensis* (e.g. various aminoglycosides, tetracyclines or chloramphenicol), we selected chloramphenicol (Cm) and gentamicin (GEN) for structural analysis. We took advantage of the fact that Cm and GEN bind the ribosome at different sites, within the peptidyl transferase center of the 50S subunit, whereas GEN binds to the decoding center of the 30S subunit, specifically within helix 44 of the 16S rRNA. We supplemented the *F. tularensis* ribosome with Cm and GEN at the same time, and collected cryo-EM data and solved the structure of the antibiotic-bound ribosome at to 2.6Å resolution (**SI Figure 8&9**).

The overall structure of the 70S ribosome - including the 30S subunit, the 50S subunit, and RaiA - was not affected by the binding of either Cm or GEN. Within this complex, Cm was found, as expected, to bind in the peptidyl transferase center of the 50S subunit (**Figure 5**). However, we identified nine molecules of GEN bound to the ribosome. As expected, one molecule occupied the decoding center of the 30S subunit, however the remaining eight molecules were not only observed within the 30S but 50S subunit as well (**Figure 6, SI Figure 10**).

Interestingly, binding of the Cm to the 50S subunit at the canonical peptidyl transferase center induced significant but localized conformational changes (**Figure 5**). A structural comparison of the 23S rRNA in the 50S subunit, with and without Cm bound, confirmed that the antibiotic physically obstructs the active site (**Figure 5b**). Detailed analysis of the 23S rRNA residues revealed the molecular impact of Cm binding. The antibiotic directly engages residues responsible for binding and catalysis, causing a rearrangement of this active site. Specifically, the Loop U2489-G2491 was found to be completely displaced upon Cm binding (**Figure 5d**). Notably, residue A2057, for which we previously did not observe any corresponding density in the 70S structure (**Figure 5c**), was clearly seen to be employed in interaction with the antibiotic in the drug-bound structure (**Figure 5d**). This demonstrates that Cm binding actively recruits this residue into the binding pocket, stabilizing the complex and locking the peptidyl transferase center into an inactive state.

### GEN binds the ribosome at multiple sites

One GEN molecule binds to the canonical decoding site, while the other copies occupy non-canonical sites (**Figure 6, SI Figure 10**). The canonical binding site for GEN is situated near the accessory protein RaiA at the 30S-50S ribosomal interface (**Figure 6b**). This site corresponds to the A-site decoding region, where aminoglycosides, such as GEN, typically interfere with cognate tRNA selection. The GEN molecule is anchored by interactions with the 16S rRNA nucleotides, including A1480-U1482, G1396-A1399. Eight additional GEN binding sites were resolved, four of them situated within the main body of the 30S subunit and five within the 50S subunit, indicating a potential mechanism for broad ribosomal perturbation beyond A-site decoding (**Figure 6**). Non-canonical Site 1 (**Figure 6c**) involves interactions with nucleobases from the 16S rRNA helix: U822-A826, U841, and U843. The second non-canonical Site 2 (**Figure 6d**) engages a distinct set of 16S rRNA residues: G1079-U1083, and A921, and C922. The remaining six molecules of GEN bind within the grooves of RNA in very similar manner with limited conformational changes (**SI Figure 10**).

## Discussion

*F. tularensis* is a potential bioterrorism agent and classified as BSL-2 or BSL-3 organism in most countries hindering its research. We aimed at the structure of its ribosome taking advantage of the existence of the BSL-2 *holarctica* strain FSC200, whose ribosome proved nearly identical to that of the notorious Schu S4 strains (**SI Figure 1**). Eventually we obtained the structure at 2.5 Å resolution.

The structure revealed a “sleeping” ribosome containing the hibernation factor RaiA. RaiA was bound to the 30S subunit in the vicinity of the 30S:50S interface, consistent with previously solved structures, and intracted almost exclusively with the 16S rRNA. Although *F. tularensis* RaiA lacks the extended C-terminal tail, we nonetheless observed only monomeric 70S ribosomes in our ribosome preparations. Although *F. tularensis* RaiA lacks the extended C-terminal tail, we nonetheless observed only monomeric 70S ribosomes in our preparations. This suggests that, even without the C-terminal tail, RaiA can outcompete the ribosome modulation factor (RMF) and the hibernation promoting factor (HPF), or that RMF/HPF were not present under the conditions that we used to grow *F. tularensis*. The RMF/HPF protein complex is known to promote the 100S ribosome dimer, whereas RaiA inhibits formation of the 100S complex [28]. The resulting 100S ribosome is resistant to degradation during stress [15, 29], which in *E. coli* is carried out primarily by RNase R and RNase II [30].

The presence of the RaiA can be attributed to the ribosome purification protocol, in which several steps involved cooling *F. tularensis* cells to 0 °C. Ribosome hibernation is a known adaptation to adverse conditions, including stress caused by low temperature [31]. However, when the same purification procedure is applied to *E. coli*, it yields ribosomes without the hibernation factor bound [32]. This observation suggests that *F. tularensis* might be very well adapted to cold. Indeed, it was observed that *F. tularensis* can remain viable for long periods at low temperatures [33, 34].

Another interesting observation was the exclusive presence of the S21 paralog 2. *F. tularensis* encodes three distinct paralogs of the small ribosomal subunit protein S21 (also known as bS21), a protein involved in translation initiation; it promotes base pairing between the 16S rRNA and the Shine-Dalgarno sequence [35, 36]. It is one of the last proteins to be incorporated during 30S subunit biogenesis [37] and can dissociate from mature ribosomes [38]. Many bacteria do not encode an S21 homolog at all, indicating that it is dispensable for protein synthesis. In *Flavobacterium johnsoniae*, depletion of S21 in the cell increases translation of reporters with strong Shine-Dalgarno sequences [39].

In our structure, we observed a clear signal corresponding to S21, suggesting high occupancy of this ribosomal subunit protein in our *F. tularensis* ribosome sample. Similarly, as in the case of RaiA, the signal was very well defined and allowed us to place side chains of amino acids residues confidently, revealing the specific presence of the paralog S21-2. In *F. tularensis*, the S21-2 paralog has been reported to be autogenously regulated [40]. Ribosomes with this paralog are uniquely important for both production of a critical virulence factor (the type VI secretion system) and intramacrophage survival, which is essential for *F. tularensis* to cause disease [41]. The preferential presence of bS21-2 might explain the observed phenomenon, that *F. tularensis* cells growing in BHI medium used in this study (mimicking the host-like environment) are more virulent than the cells cultivated in non-host-like environments [42]. However, why this paralog of bS21 protein is preferentially present in the hibernating ribosomes of *F. tularensis* remains to be established.

Because the ribosome is a major target of antibiotics and *F. tularensis* is a dangerous pathogen with bioterrorism potential, we determined the structure of its ribosome bound to two antibiotics, chloramphenicol (Cm) and gentamicin (GEN). Remarkably, comparison of the Cm-bound ribosome with the unliganded ribosome revealed that the peptidyl-transferase center (PTC) adopts an unusual conformation in the unliganded state. Cm binding, however, induces a substantial rearrangement of the PTC in the 50S subunit, shifting it toward a configuration more similar to that seen in other bacterial ribosomes [43, 44].

Interestingly, in the case of GEN, in addition to the canonical decoding-center site, we detected eight further GEN molecules scattered across both ribosomal subunits, each engaging distinct rRNA pockets. However, this is not unprecedented in the literature, structures with multiple GEN molecules bound to the ribosome were reported before [45–49]. These additional, non-canonical binding modes suggest a broader, multifaceted mechanism of ribosomal disruption. Nevertheless, the primary mechanism of GEN function, similarly to other bacteria, is also in *F. tularensis* the inhibition of the A-site as supported by our structural analysis.

Together, our findings provide the structural understanding of the *F. tularensis* ribosome. By revealing *F. tularensis*-specific structural features and both canonical and non-canonical binding pockets, our work presents opportunities for rational antibiotic design. It also provides mechanistic understanding of ribosome hibernation, most likely an adaptation of *F. tularensis* to moist environments like water, soil, hay, straw or decaying animal carcasses [33, 34].

## Materials and Methods

### Ribosome purification

The ribosomes were prepared according to Cui et al. (2022) with minor modifications [32]. The *Francisella tularensis* subsp. *holarctica* strain FSC200 was inoculated onto McLeod agar plates and incubated overnight (ON) at 37°C. Colonies from the McLeod plate were transferred into 20 mL brain heart infusion (BHI) medium (BD, 211059) supplemented with 0.1% (w/v) cysteine (Sigma-Aldrich, 30120) and cultured under aerobic conditions at 37°C overnight. The overnight culture was thereafter inoculated into 400 mL of fresh BHI medium (enriched by 0.1% cysteine) and incubated for approximately 2 hours under the same conditions. Subsequently, the culture was scaled up to 2 L BHI medium (enriched by 0.1% cysteine) and grown until the optical density at 600 nm (OD_600_) reached 0.8–1.0.

The bacterial culture was gradually cooled from 37 °C to 4 °C to facilitate run-off ribosome production, then centrifuged at 4,000 rpm for 30 min at 4°C. The cell pellet was washed three times with ice-cold phosphate-buffered saline (PBS) with subsequent centrifugation (using Beckman SW41 Ti rotor) at 10000 rpm for 10 min at 4 °C. Residual PBS was carefully removed, and the pellet was weighed. For lysis, S30A buffer (14 mM L-glutamic acid hemimagnesium salt tetrahydrate, 60 mM potassium L-glutamate, 50 mM Tris, pH 7.7) was added at 1 ml per gram of cell pellet, and cells were resuspended thoroughly. The suspension was lysed by three passes through a French pressure cell press (Thermo Fisher Scientific) at 16,000 psi. The lysate was incubated at 37°C and 220 rpm for 80 min to allow ribosome run-off. Following digestion, the sample was centrifuged at 10,000 rpm for 60 min at 4 °C, and the supernatant was collected and centrifuged again under the same conditions.

The clarified lysate was filtered through a 0.22 μm sterile membrane. Equal volumes of the filtrate and centrifugation buffer (20 mM Tris pH 7.7, 500 mM ammonium chloride, 20 mM magnesium acetate, 0.5 mM EDTA, 7 mM β-mercaptoethanol, 30% sucrose) were combined in ultracentrifuge tubes and centrifuged at 170,000 × g for 2 h at 4 °C. The supernatant was discarded, and the pellet was resuspended in precooled S30A buffer on ice. The resuspension was again mixed 1:1 with ultracentrifugation buffer and subjected to a second ultracentrifugation step at 170,000×g for 2 hours at 4°C. The resulting pellet was resuspended in 500 μl precooled ribosome buffer (20 mM HEPES pH 7.7, 20 mM magnesium acetate, 30 mM potassium chloride, 7 mM β-mercaptoethanol) and transferred to sterile 1.5 ml tubes.

The protein concentration was estimated to be 6.2 mg/mL using the Qubit method (Thermo Fisher Scientific, A50668). Finally, the quality of the ribosomes was checked using agarose and SDS-PAGE electrophoresis (SI Figure 1) and the ribosomes were aliquoted and stored at -80°C until needed.

#### Cryo-EM Sample Preparation and Data Processing

Purified *F. tularensis* 70S ribosomes, diluted to ∼ 20 μM in buffer containing 20 mM HEPES (pH 7.5), 30 mM KCl, 20 mM MgCl₂, and 7 mM β-mercaptoethanol. Aliquots (3 μL) were applied onto glow-discharged Quantifoil R2/1 300-mesh copper grids (Electron Microscopy Sciences, Prod. No. Q350CR1) that had been glow discharged at 15 mA for 30 s immediately prior to vitrification. Grids were blotted for 5 s with a 5 s wait time and blot force of –5 at 4°C and 100 % humidity, then plunge-frozen in liquid ethane using a Vitrobot Mark IV (Thermo Fisher Scientific).

Cryo-EM data were collected on a Titan Krios electron microscope operated at 300 kV and equipped with a direct electron detector. A total of 28,005 movies were recorded at a pixel size of 0.76 Å with a total electron dose of 40 e⁻/Å². After initial curation, 24,869 high-quality movies were selected for further processing. Motion correction and contrast transfer function (CTF) estimation were performed in CryoSPARC v4.6.2. An initial subset of 160 movies was processed, from which 16,203 particles were selected for 2D classification. Representative 2D templates generated from this dataset were then used for template-based particle picking across the curated 24,869 movies, yielding 6,845,738 particles. Following particle curation, 2,039,854 particles were selected and extracted (box size 600 pixels, Fourier-cropped to 150 pixels) and subjected to multiple rounds of 2D classification.

From these, 872,346 particles were retained for *ab initio* 3D reconstruction, which produced three initial classes. Class 0 (104,870 particles), Class 1 (291,093 particles), and Class 2 (476,383 particles) were refined independently. The best-resolved class (Class 2) was subjected to homogeneous and 3D refinement, yielding the final reconstruction. The overall resolution of the *F. tularensis* 70S ribosome map was 2.5 Å, as determined by the gold-standard Fourier shell correlation (GSFSC) 0.143 criterion.

The second *F. tularensis* 70S dataset was processed similarly and collected under identical conditions. Briefly, the solution of the 70S ribosome was supplied with ∼100 fold molar excess of the antibiotics chloramphenicol and gentamicin and incubated on ice for 2 hours. Grids were prepared, frozen, and the dataset was collected as described above. Processing of the antibiotic-bound 70S dataset followed an analogous workflow with only minor adjustments (see Supplementary Information).

In the final reconstruction, particles from 5,468 movies were used after motion and CTF correction. Of these, 181 movies were used as a subset to obtain initial 2D classes for reference-based template picking and for generating the initial model. The final cryo-EM map was reconstructed from 149,692 particles, reaching an estimated resolution of 2.6 Å according to the gold-standard Fourier shell correlation (GSFSC) 0.143 criterion. The final maps and the refined model were deposited under PDB ID 9T6H.

#### Model building and refinement

Initially, the template 70S ribosome structure of *Staphylococcus aureus* (pdb entry 5LI0) [20] was fitted to density using ChimeraX [50]. The 23S, 16S, and 5S ribosomal RNA sequences of *Francisella tularensis* subsp. *holarctica* FSC200 were extracted from the GenBank entry NC_019551.1, aligned with the *S. aureus* rRNA sequences using ApE v3.1.3 [51] and the rRNA molecules in the template model were accordingly mutated using Coot v0.9.8.7 [52]. Next, structures of the individual ribosomal proteins of *F. tularensis* predicted by AlphaFold v2.0 [53, 54] were aligned with the template model using the LSQ and SSM Superpose algorithms in Coot. AlphaFold models of the 30S ribosomal protein S21 variant 2 and the ribosome-associated translation inhibitor (ribosome hibernation promoting factor) RaiA were fitted to density manually in Coot. The model was further improved to good geometry and correlation coefficients using automatic model refinement with the phenix.real_space_refine tool from the Phenix package v1.20.1-4487 [55] and manual model building with Coot. Geometrical restraints for N2-methylguanosine-5’-phosphate were generated with Grade2 v1.3.1 (Global Phasing Ltd.). The atomic coordinates, structural factors and cryo-EM were deposited in the Protein Data Bank (https://www.rcsb.org) under the accession code 9SDA.

For antibiotics containing 70S ribosome. The final model was build using aformention 70S structure (9SDA) as a starting model. Additionally one molecule of chloramphenicol and nine molecules of gentamicin were convincingly fitted into the cryo-EM density. The final maps and the model of 70S with antibiotics were deposited under accession code 9T6H in the Protein Data Bank (https://www.rcsb.org). In parallel all maps including half-maps for both structures were co-deposited to EM Data Bank under accession codes EMDB-54782, and EMDB-55615 respectively. The collection and final refinement statistics are shown in Supplementary Table 1.

## Acknowledgments

The research was funded by the project”New Technologies for Translational Research in Pharmaceutical Sciences” /NETPHARM, project ID CZ.02.01.01/00/22_008/0004607 (co-funded by the European Union). We are greatful for help with cryo-EM data collection to the IOCB cryo-EM facility and the CIISB cryo-EM facility, Instruct-CZ Centre, supported by MEYS CR (LM2023042) and European Regional Development Fund-Project „Innovation of Czech Infrastructure for Integrative Structural Biology“ (No. CZ.02.01.01/00/23_015/0008175).

## Contributions

M.K, J.S., and P.P.,performed experiments. M.K. and J.S. analyzed the cryo-EM data. K.H. conceived the project. All authors co-wrote the manuscript.

