## Supplementary Information for "Structure of the Hibernating *Francisella tularensis* Ribosome and Mechanistic Insights into Its Inhibition by Antibiotics"

a

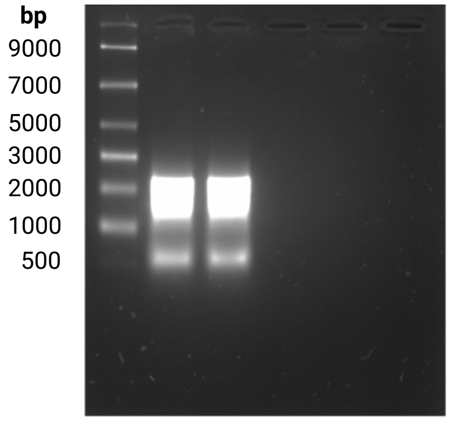

b

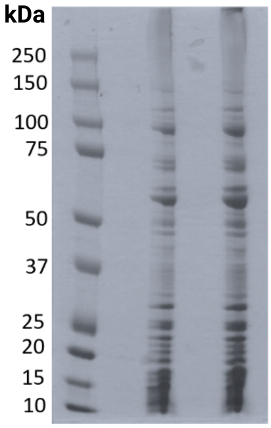

c

| **Alignment results link** | | [CLICK HERE]() | | | | | | |
|---|---|---|---|---|---|---|---|---|
| **Entry** | **Reviewed** | **Entry Name** | **Protein names** | **Gene Names** | **Organism** | **Length** | **Identity (Pairwise Sequence Alignment) SCHU S4 vs FSC200 (%)** | **Similarity (Pairwise Sequence Alignment) SCHU S4 vs FSC200 (%)** |
| Q5NID3 | reviewed | RL7_FRATT | Large ribosomal subunit protein bL12 (50S ribosomal protein L7/L12) | rplL FTT_0143 | Francisella tularensis subsp. tularensis (strain SCHU S4 / Schu 4) | 125 | 98,4 | 98,4 |
| Q5NHU2 | reviewed | RL17_FRATT | Large ribosomal subunit protein bL17 (50S ribosomal protein L17) | rplQ FTT_0351 | Francisella tularensis subsp. tularensis (strain SCHU S4 / Schu 4) | 145 | 99,3 | 99,3 |
| Q5NIC3 | reviewed | RL19_FRATT | Large ribosomal subunit protein bL19 (50S ribosomal protein L19) | rplS FTT_0153 | Francisella tularensis subsp. tularensis (strain SCHU S4 / Schu 4) | 115 | 98,3 | 99,1 |
| Q5NGL5 | reviewed | RL20_FRATT | Large ribosomal subunit protein bL20 (50S ribosomal protein L20) | rplT FTT_0820 | Francisella tularensis subsp. tularensis (strain SCHU S4 / Schu 4) | 118 | 98,3 | 98,3 |
| Q5NGR2 | reviewed | RL21_FRATT | Large ribosomal subunit protein bL21 (50S ribosomal protein L21) | rplU FTT_0772 | Francisella tularensis subsp. tularensis (strain SCHU S4 / Schu 4) | 104 | 99 | 99 |
| Q5NH01 | reviewed | RL25_FRATT | Large ribosomal subunit protein bL25 (50S ribosomal protein L25) | rplY FTT_0675 | Francisella tularensis subsp. tularensis (strain SCHU S4 / Schu 4) | 96 | 99 | 99 |
| Q5NGR1 | reviewed | RL27_FRATT | Large ribosomal subunit protein bL27 (50S ribosomal protein L27) | rpmA FTT_0773 | Francisella tularensis subsp. tularensis (strain SCHU S4 / Schu 4) | 84 | 98,8 | 98,8 |
| Q5NEM3 | reviewed | RL28_FRATT | Large ribosomal subunit protein bL28 (50S ribosomal protein L28) | rpmB FTT_1603 | Francisella tularensis subsp. tularensis (strain SCHU S4 / Schu 4) | 78 | 100 | 100 |

| | | | | | | | | |
|---|---|---|---|---|---|---|---|---|
| Q5NHS9 | reviewed | RL31_FRATT | Large ribosomal subunit protein bL31 (50S ribosomal protein L31) | rpmE FTT_0366 | Francisella tularensis subsp. tularensis (strain SCHU S4 / Schu 4) | 71 | 97,2 | 97,2 |
| Q5NF72 | reviewed | RL32_FRATT | Large ribosomal subunit protein bL32 (50S ribosomal protein L32) | rpmF FTT_1371 | Francisella tularensis subsp. tularensis (strain SCHU S4 / Schu 4) | 60 | 98,3 | 100 |
| Q5NEM2 | reviewed | RL33_FRATT | Large ribosomal subunit protein bL33 (50S ribosomal protein L33) | rpmG FTT_1604 | Francisella tularensis subsp. tularensis (strain SCHU S4 / Schu 4) | 51 | 100 | 100 |
| Q5NI53 | reviewed | RL34_FRATT | Large ribosomal subunit protein bL34 (50S ribosomal protein L34) | rpmH FTT_0236c | Francisella tularensis subsp. tularensis (strain SCHU S4 / Schu 4) | 44 | 100 | 100 |
| Q5NGL6 | reviewed | RL35_FRATT | Large ribosomal subunit protein bL35 (50S ribosomal protein L35) | rpmI FTT_0819 | Francisella tularensis subsp. tularensis (strain SCHU S4 / Schu 4) | 65 | 100 | 100 |
| Q5NHU7 | reviewed | RL36_FRATT | Large ribosomal subunit protein bL36 (50S ribosomal protein L36) | rpmJ FTT_0346 | Francisella tularensis subsp. tularensis (strain SCHU S4 / Schu 4) | 37 | 100 | 100 |
| Q5NG01 | reviewed | RL9_FRATT | Large ribosomal subunit protein bL9 (50S ribosomal protein L9) | rplI FTT_1060c | Francisella tularensis subsp. tularensis (strain SCHU S4 / Schu 4) | 151 | 99,3 | 99,3 |
| Q5NID5 | reviewed | RL1_FRATT | Large ribosomal subunit protein uL1 (50S ribosomal protein L1) | rplA FTT_0141 | Francisella tularensis subsp. tularensis (strain SCHU S4 / Schu 4) | 231 | 99,6 | 100 |
| Q5NID4 | reviewed | RL10_FRATT | Large ribosomal subunit protein uL10 (50S ribosomal protein L10) | rplJ FTT_0142 | Francisella tularensis subsp. tularensis (strain SCHU S4 / Schu 4) | 172 | 98,8 | 100 |
| Q5NID6 | reviewed | RL11_FRATT | Large ribosomal subunit protein uL11 (50S ribosomal protein L11) | rplK FTT_0140 | Francisella tularensis subsp. tularensis (strain SCHU S4 / Schu 4) | 144 | 99,3 | 100 |
| Q5NFG3 | unreviewed | Q5NFG3_FRATT | Large ribosomal subunit protein uL13 | rplM FTT_1273 | Francisella tularensis subsp. tularensis (strain SCHU S4 / Schu 4) | 151 | 94 | 94 |
| Q5NHV8 | reviewed | RL14_FRATT | Large ribosomal subunit protein uL14 (50S ribosomal protein L14) | rplN FTT_0335 | Francisella tularensis subsp. tularensis (strain SCHU S4 / Schu 4) | 122 | 100 | 100 |
| Q5NHU9 | reviewed | RL15_FRATT | Large ribosomal subunit protein uL15 (50S ribosomal protein L15) | rplO FTT_0344 | Francisella tularensis subsp. tularensis (strain SCHU S4 / Schu 4) | 143 | 98,6 | 98,6 |
| Q5NHW1 | reviewed | RL16_FRATT | Large ribosomal subunit protein uL16 (50S ribosomal protein L16) | rplP FTT_0332 | Francisella tularensis subsp. tularensis (strain SCHU S4 / Schu 4) | 137 | 99,3 | 100 |
| Q5NHV2 | reviewed | RL18_FRATT | Large ribosomal subunit protein uL18 (50S | rplR FTT_0341 | Francisella tularensis subsp. tularensis (strain | 117 | 100 | 100 |

| | | | | | | | | |
|---|---|---|---|---|---|---|---|---|
| | | | ribosomal protein L18) | | SCHU S4 / Schu 4) | | | |
| Q5NHW5 | reviewed | RL2_FRATT | Large ribosomal subunit protein uL2 (50S ribosomal protein L2) | rplB FTT_0328 | Francisella tularensis subsp. tularensis (strain SCHU S4 / Schu 4) | 274 | 99,6 | 100 |
| Q5NHW3 | reviewed | RL22_FRATT | Large ribosomal subunit protein uL22 (50S ribosomal protein L22) | rplV FTT_0330 | Francisella tularensis subsp. tularensis (strain SCHU S4 / Schu 4) | 111 | 100 | 100 |
| Q5NHW6 | reviewed | RL23_FRATT | Large ribosomal subunit protein uL23 (50S ribosomal protein L23) | rplW FTT_0327 | Francisella tularensis subsp. tularensis (strain SCHU S4 / Schu 4) | 99 | 99 | 99 |
| Q5NHV7 | reviewed | RL24_FRATT | Large ribosomal subunit protein uL24 (50S ribosomal protein L24) | rplX FTT_0336 | Francisella tularensis subsp. tularensis (strain SCHU S4 / Schu 4) | 105 | 99 | 99 |
| Q5NHW0 | reviewed | RL29_FRATT | Large ribosomal subunit protein uL29 (50S ribosomal protein L29) | rpmC FTT_0333 | Francisella tularensis subsp. tularensis (strain SCHU S4 / Schu 4) | 66 | 98,5 | 98,5 |
| Q5NHW8 | reviewed | RL3_FRATT | Large ribosomal subunit protein uL3 (50S ribosomal protein L3) | rplC FTT_0325 | Francisella tularensis subsp. tularensis (strain SCHU S4 / Schu 4) | 211 | 99,5 | 99,5 |
| Q5NHV0 | reviewed | RL30_FRATT | Large ribosomal subunit protein uL30 (50S ribosomal protein L30) | rpmD FTT_0343 | Francisella tularensis subsp. tularensis (strain SCHU S4 / Schu 4) | 61 | 100 | 100 |
| Q5NHW7 | reviewed | RL4_FRATT | Large ribosomal subunit protein uL4 (50S ribosomal protein L4) | rplD FTT_0326 | Francisella tularensis subsp. tularensis (strain SCHU S4 / Schu 4) | 207 | 100 | 100 |
| Q5NHV6 | reviewed | RL5_FRATT | Large ribosomal subunit protein uL5 (50S ribosomal protein L5) | rplE FTT_0337 | Francisella tularensis subsp. tularensis (strain SCHU S4 / Schu 4) | 179 | 99,4 | 99,4 |
| Q5NHV3 | reviewed | RL6_FRATT | Large ribosomal subunit protein uL6 (50S ribosomal protein L6) | rplF FTT_0340 | Francisella tularensis subsp. tularensis (strain SCHU S4 / Schu 4) | 178 | 99,4 | 100 |
| Q5NI98 | unreviewed | Q5NI98_FRATT | Small ribosomal subunit protein bS1 (30S ribosomal protein S1) | rpsA FTT_0183c | Francisella tularensis subsp. tularensis (strain SCHU S4 / Schu 4) | 556 | 99,5 | 99,6 |
| Q5NIC6 | reviewed | RS16_FRATT | Small ribosomal subunit protein bS16 (30S ribosomal protein S16) | rpsP FTT_0150 | Francisella tularensis subsp. tularensis (strain SCHU S4 / Schu 4) | 82 | 98,8 | 98,8 |
| Q5NG00 | reviewed | RS18_FRATT | Small ribosomal subunit protein bS18 (30S ribosomal protein S18) | rpsR FTT_1061c | Francisella tularensis subsp. tularensis (strain SCHU S4 / Schu 4) | 72 | 97,2 | 98,6 |
| Q5NEF7 | reviewed | RS20_FRATT | Small ribosomal subunit protein bS20 (30S ribosomal protein S20) | rpsT FTT_1679 | Francisella tularensis subsp. tularensis (strain SCHU S4 / Schu 4) | 90 | 98,9 | 98,9 |
| Q5NHQ | reviewed | RS211_FRAT | Small ribosomal | rpsU1 | Francisella | 65 | 96,9 | 98,5 |

| | | | | | | | | |
|---|---|---|---|---|---|---|---|---|
| 7 | | T | subunit protein bS21A (30S ribosomal protein S21 1) | FTT_0390c | tularensis subsp. tularensis (strain SCHU S4 / Schu 4) | | | |
| Q5NGS8 | reviewed | RS212_FRATT | Small ribosomal subunit protein bS21B (30S ribosomal protein S21 2) | rpsU2 FTT_0753 | Francisella tularensis subsp. tularensis (strain SCHU S4 / Schu 4) | 66 | 100 | 100 |
| Q5NG21 | reviewed | RS213_FRATT | Small ribosomal subunit protein bS21C (30S ribosomal protein S21 3) | rpsU3 FTT_1038c | Francisella tularensis subsp. tularensis (strain SCHU S4 / Schu 4) | 65 | 100 | 100 |
| Q5NFZ9 | reviewed | RS6_FRATT | Small ribosomal subunit protein bS6 (30S ribosomal protein S6) | rpsF FTT_1062c | Francisella tularensis subsp. tularensis (strain SCHU S4 / Schu 4) | 111 | 100 | 100 |
| Q5NHW9 | reviewed | RS10_FRATT | Small ribosomal subunit protein uS10 (30S ribosomal protein S10) | rpsJ FTT_0324 | Francisella tularensis subsp. tularensis (strain SCHU S4 / Schu 4) | 105 | 99 | 100 |
| Q5NHU5 | reviewed | RS11_FRATT | Small ribosomal subunit protein uS11 (30S ribosomal protein S11) | rpsK FTT_0348 | Francisella tularensis subsp. tularensis (strain SCHU S4 / Schu 4) | 129 | 98,4 | 99,2 |
| Q5NHX2 | reviewed | RS12_FRATT | Small ribosomal subunit protein uS12 (30S ribosomal protein S12) | rpsL FTT_0321 | Francisella tularensis subsp. tularensis (strain SCHU S4 / Schu 4) | 124 | 100 | 100 |
| Q5NHU6 | reviewed | RS13_FRATT | Small ribosomal subunit protein uS13 (30S ribosomal protein S13) | rpsM FTT_0347 | Francisella tularensis subsp. tularensis (strain SCHU S4 / Schu 4) | 118 | 100 | 100 |
| Q5NHV5 | reviewed | RS14_FRATT | Small ribosomal subunit protein uS14 (30S ribosomal protein S14) | rpsN FTT_0338 | Francisella tularensis subsp. tularensis (strain SCHU S4 / Schu 4) | 101 | 100 | 100 |
| Q5NGX8 | reviewed | RS15_FRATT | Small ribosomal subunit protein uS15 (30S ribosomal protein S15) | rpsO FTT_0698 | Francisella tularensis subsp. tularensis (strain SCHU S4 / Schu 4) | 88 | 98,9 | 100 |
| Q5NHV9 | reviewed | RS17_FRATT | Small ribosomal subunit protein uS17 (30S ribosomal protein S17) | rpsQ FTT_0334 | Francisella tularensis subsp. tularensis (strain SCHU S4 / Schu 4) | 83 | 98,8 | 98,8 |
| Q5NHW4 | reviewed | RS19_FRATT | Small ribosomal subunit protein uS19 (30S ribosomal protein S19) | rpsS FTT_0329 | Francisella tularensis subsp. tularensis (strain SCHU S4 / Schu 4) | 92 | 100 | 100 |
| Q5NHY0 | reviewed | RS2_FRATT | Small ribosomal subunit protein uS2 (30S ribosomal protein S2) | rpsB FTT_0313 | Francisella tularensis subsp. tularensis (strain SCHU S4 / Schu 4) | 239 | 98,7 | 98,7 |
| Q5NHW2 | reviewed | RS3_FRATT | Small ribosomal subunit protein uS3 (30S ribosomal protein S3) | rpsC FTT_0331 | Francisella tularensis subsp. tularensis (strain SCHU S4 / Schu 4) | 222 | 98,2 | 98,7 |
| Q5NHU4 | reviewed | RS4_FRATT | Small ribosomal subunit protein uS4 | rpsD FTT_0349 | Francisella tularensis subsp. tularensis (strain SCHU S4 / Schu | 206 | 100 | 100 |

| | | | | | | | | |
|---|---|---|---|---|---|---|---|---|
| | | | | | 4) | | | |
| Q5NHV1 | reviewed | RS5_FRATT | Small ribosomal subunit protein uS5 (30S ribosomal protein S5) | rpsE FTT_0342 | Francisella tularensis subsp. tularensis (strain SCHU S4 / Schu 4) | 166 | 99,4 | 100 |
| Q5NHX1 | reviewed | RS7_FRATT | Small ribosomal subunit protein uS7 (30S ribosomal protein S7) | rpsG FTT_0322 | Francisella tularensis subsp. tularensis (strain SCHU S4 / Schu 4) | 157 | 76,4 | 76,4 |
| Q5NHV4 | reviewed | RS8_FRATT | Small ribosomal subunit protein uS8 (30S ribosomal protein S8) | rpsH FTT_0339 | Francisella tularensis subsp. tularensis (strain SCHU S4 / Schu 4) | 132 | 99,2 | 100 |
| Q5NFG2 | unreviewed | Q5NFG2_FRATT | Small ribosomal subunit protein uS9 (30S ribosomal protein S9) | rpsI FTT_1274 | Francisella tularensis subsp. tularensis (strain SCHU S4 / Schu 4) | 132 | 100 | 100 |
| | | | **Average percentage of identity/similarity** | | | | **98,71785714** | **98,975** |
| **rRNA alignment** | | | | | | | | |
| **Locus tag** | **Locus tag** | **Locus tag** | **Locus tag** | **Gene** | **Strain** | | **Identity (Pairwise Sequence Alignment) SCHU S4 vs FSC200 (%)** | **Similarity (Pairwise Sequence Alignment) SCHU S4 vs FSC200 (%)** |
| FTT_r10 | FTT_r04 | FTT_r07 | | 16S rRNA | Francisella tularensis subsp. tularensis (strain SCHU S4 / Schu 4) | | 99,3 | 99,3 |
| FTS_1142 | FTS_0441 | FTS_0114 | | 16S rRNA | Francisella tularensis subsp. holarctica (strain FSC200) | | | |
| FTT_r09 | FTT_r06 | FTT_r03 | | 23S rRNA | Francisella tularensis subsp. tularensis (strain SCHU S4 / Schu 4) | | 98,7 | 98,7 |
| FTS_1145 | FTS_0444 | FTS_0117 | | 23S rRNA | Francisella tularensis subsp. holarctica (strain FSC200) | | | |
| FTT_r01 | FTT_r02 | FTT_r05 | FTT_r08 | 5S rRNA | Francisella tularensis subsp. tularensis (strain SCHU S4 / Schu 4) | | 95,7 | 95,7 |
| FTS_1146 | FTS_0475 | FTS_0445 | FTS_0118 | 5S rRNA | Francisella tularensis subsp. holarctica (strain FSC200) | | | |
| | | | **Average percentage of identity/similarity** | | | | **97,9** | **97,9** |

**SI Figure 1. RNA electrophoresis, SDS-PAGE, and Pairwise alignment of ribosomal subunits**
**a** RNA electrophoresis. RNA content in ribosome samples was analyzed by agarose gel electrophoresis. Samples were mixed with RNA loading dye (New England Biolabs) and loaded onto a 1% agarose gel. Electrophoresis was performed at 100 V for 30 min, and RNA bands were visualized under UV illumination. 1. line – ladder, 2. and 3. lines – two replicates of ribosome samples
**b** Protein electrophoresis. Protein content in ribosome preparations was analyzed by SDS–polyacrylamide gel electrophoresis (SDS–PAGE). Samples were loaded onto a 12% polyacrylamide gel and electrophoresed at 200 V for 45 min. 1. line – ladder, 2. and 3. lines – two replicates of ribosome samples
**c** Pairwise alignment of ribosomal subunits FSC200 vs. Schu S4. Each subunit of FSC200 and Schu S4 ribosome was aligned using pairwise sequence alignment tool available online at: www.ebi.ac.uk/jdispatcher/psa/emboss_needle

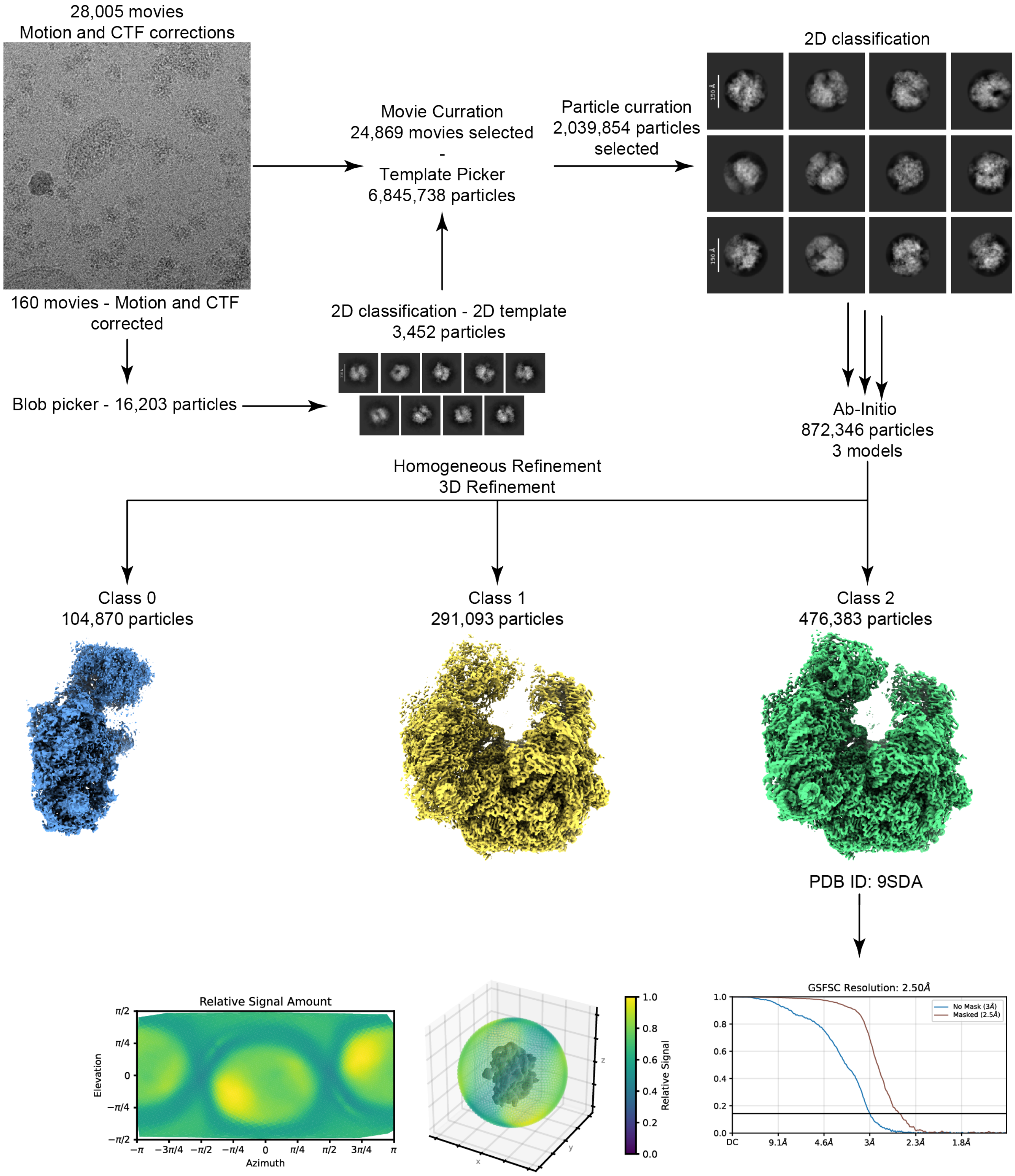

**SI Figure 2. Workflow *F. tularensis* 70S**
Cryo-EM processing from raw movies collected on Titan Krios with Falcon 4i detector containing sample with *F. tularensis* 70S. Straightforward cryosparc processing lead to final 3D reconstruction with GSFSC final resolution at 2.5 Å.

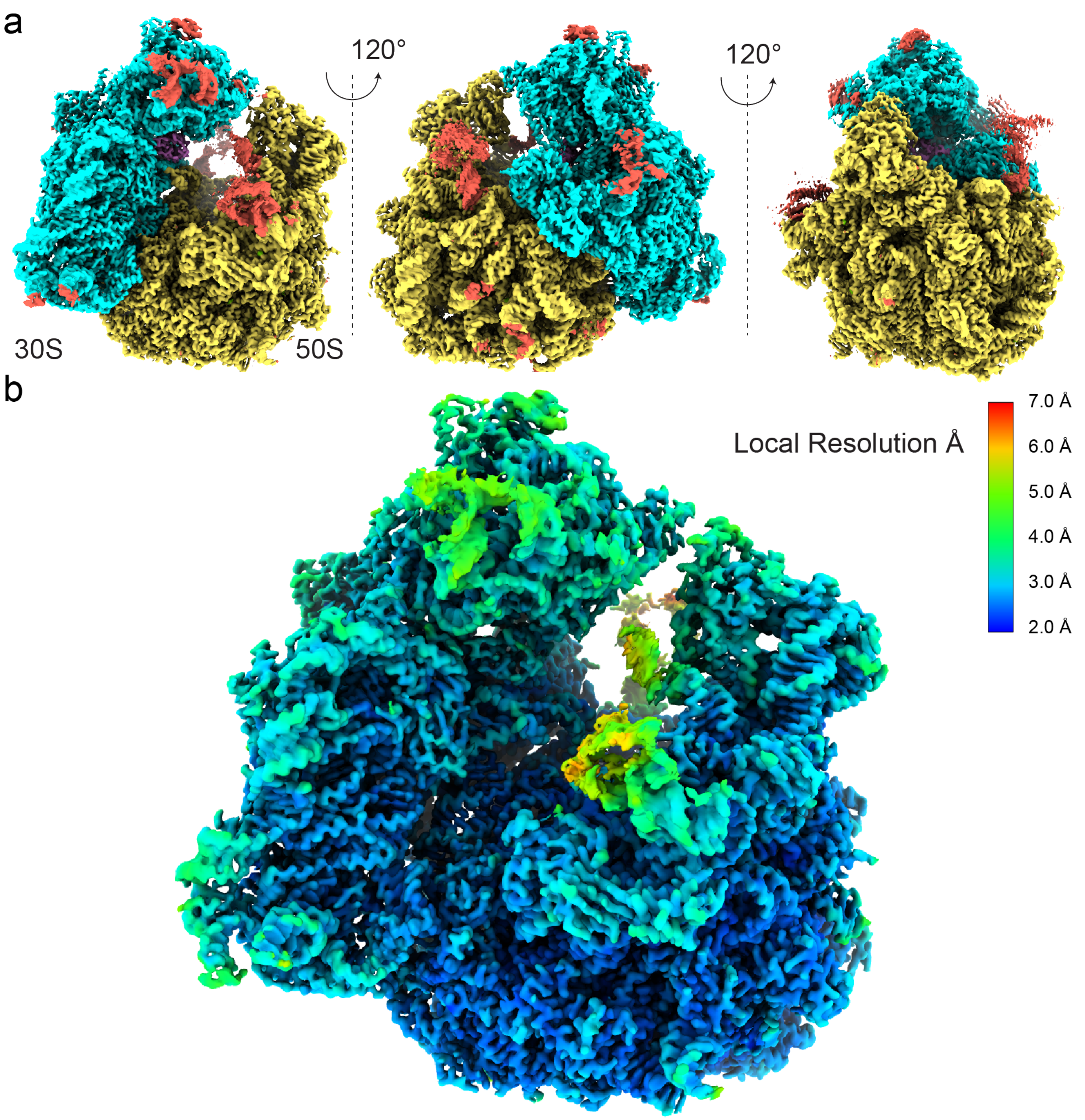

**SI Figure 3. CryoEM maps of 70S ribosome**

**a** Cryo-EM maps of *F. tularensis* 70S ribosome coloured by subunits; 30S cyan, 50S yellow, and RaiA violet. Maps with poor density that did not allow model building are shown in red.

**b** Cryo-EM map coloured by local resolution estimation.

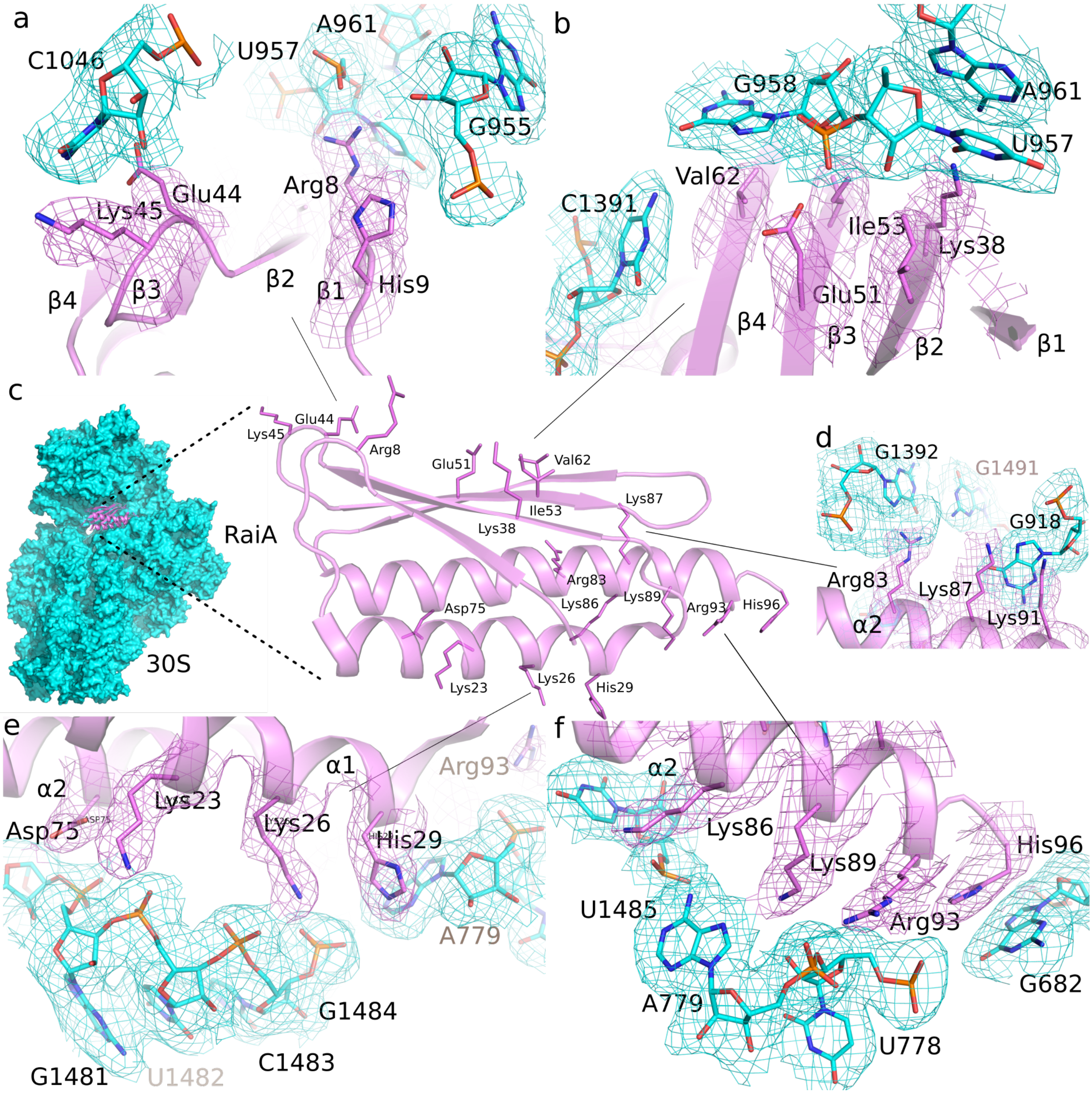

**SI Figure 4. Cryo-EM maps**

Cryo-EM map depicting the vicinity of interaction between RaiA (violet) and 30S of *F. tularensis* (cyan). For clarity, the layout is identical to Figure 2. The residues within the of interaction distance 3.6 A are shown as well as cryo-EM map in their vicinity, coloured according to the residues (maps are contoured to sigma = 2.0).

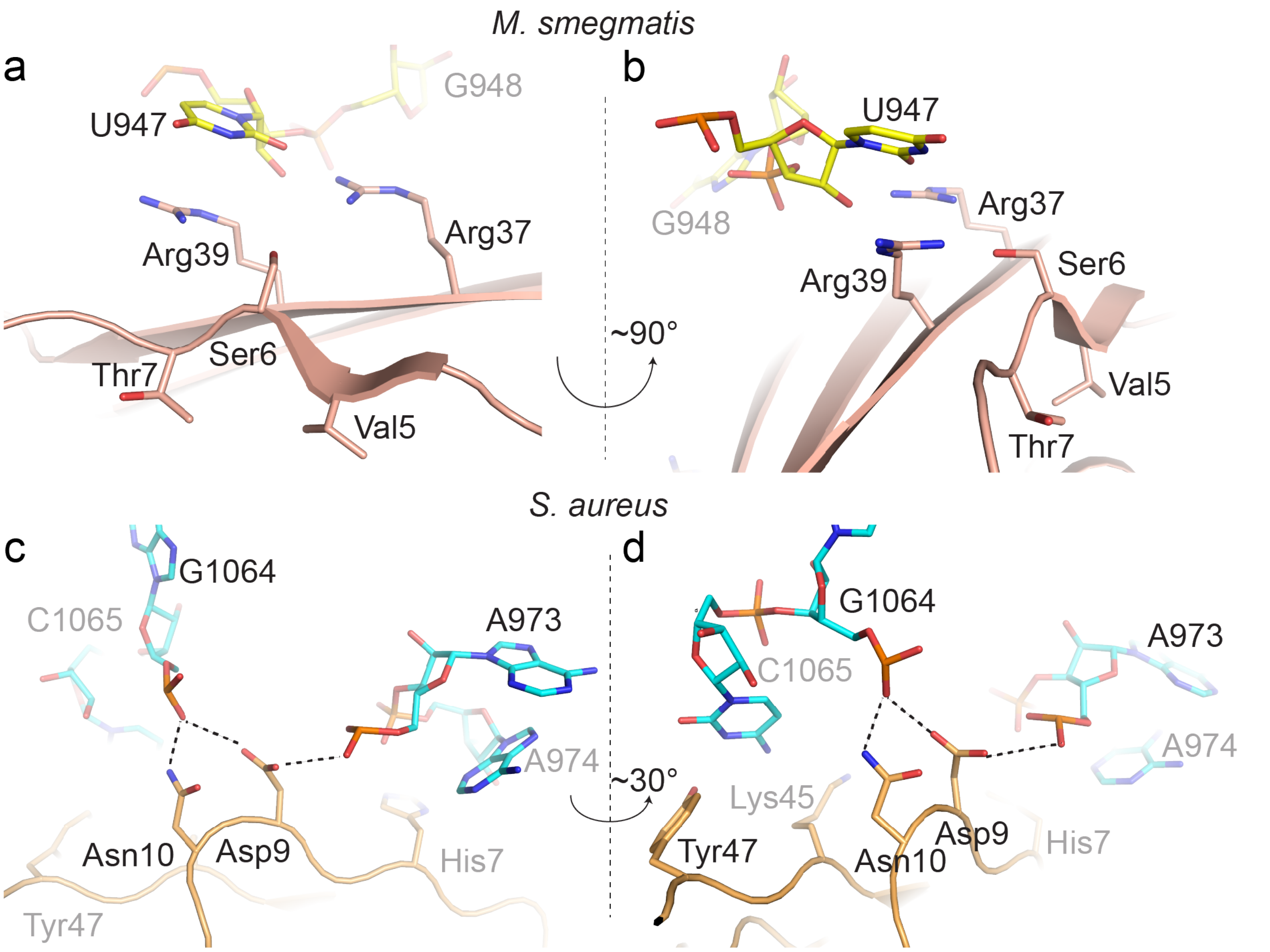

**SI Figure 5. Details of interaction between *F. tularensis* RaiA homologous proteins within the 70S from different bacteria.**
(a) Detail of interaction of Arg39 from RafH and U947 from 16S rRNA from *M. smegmatis* (PDBID 8WHX). (b) Details of interaction hibernation promoting factor HPF/YfiA family from *Staphylococcus aureus* between Asp9 and Asn10 with 16S rRNA (PDBID 6S0X).

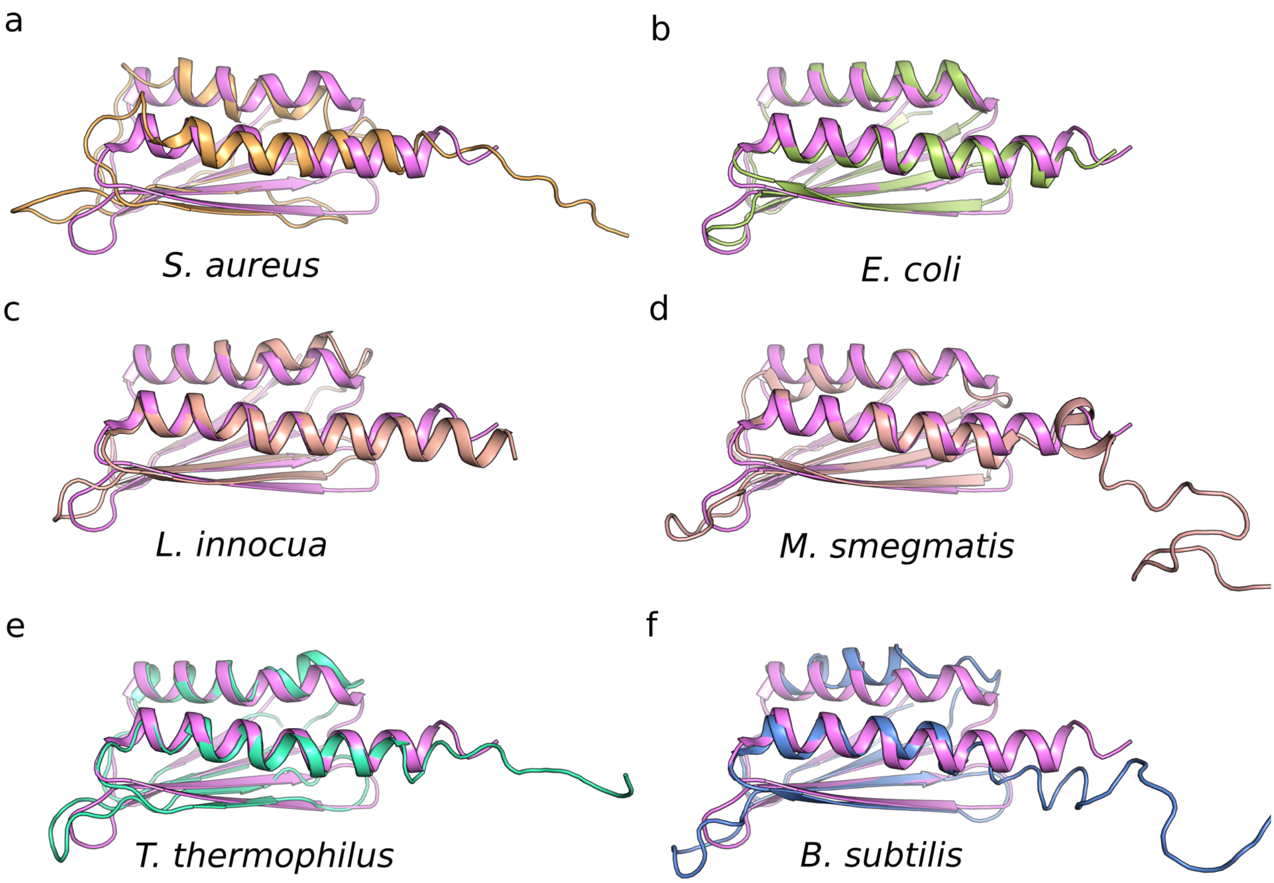

**SI Figure 6. Structural alignment of *F. tularensis* RaiA with differences in Ribosome Hiberenation Factors among representative species**
alignment *of F. tularensis RaiA* with **a** *S. aureus*, **b** *E. coli*, **c** *Listeria innocua*, **d** *M. smegmatis* RafH, **e** *B. subtilis*, **f** *T. thermophilus.*

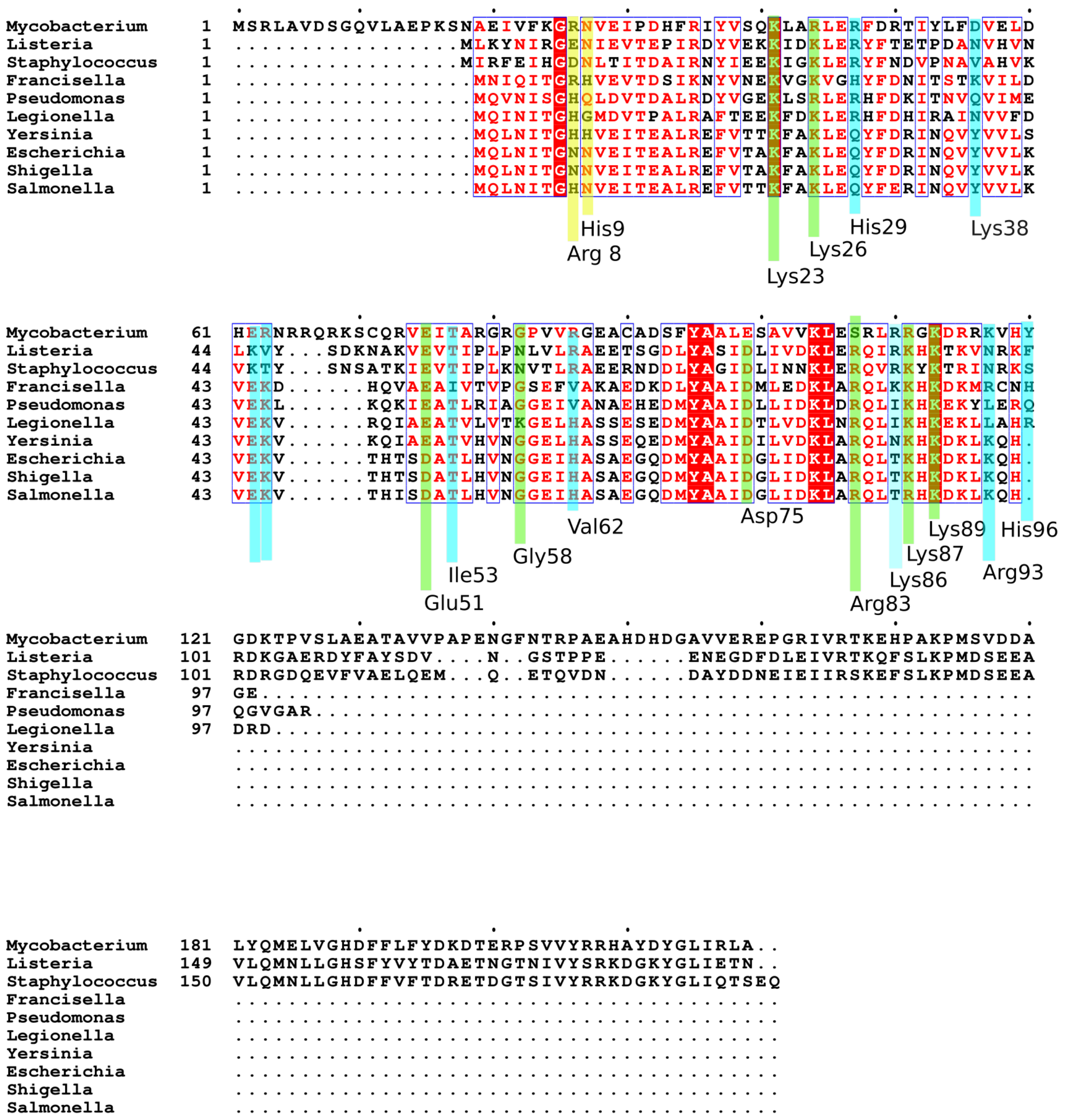

**SI Figure 7. RaiA alignments**

Sequence alignment of *F. tularensis* RaiA with homologous proteins from selected species. The residues responsible for interaction with 16S rRNA of 30S subunit are highlighted (green for high degree of homology, cyan and yellow for lower degrees of homology).

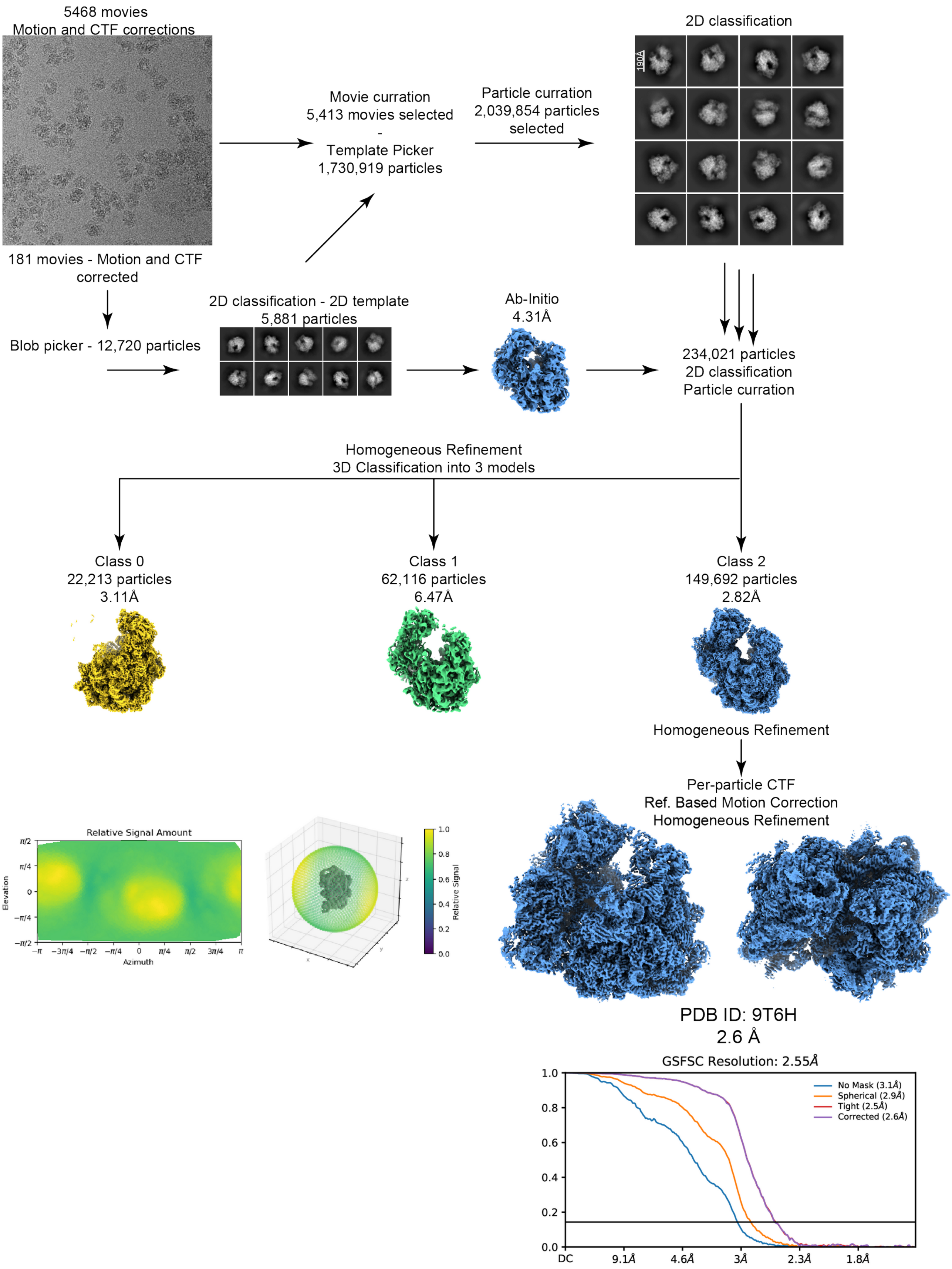

**SI Figure 8. Workflow *F. tularensis* 70S with Antibiotics Cm&GEN**
Cryo-EM processing of *F. tularensis* 70S with antibiotics chloramphenicol and gentamicin including the final 3D reconstruction with GSFSC final resolution.

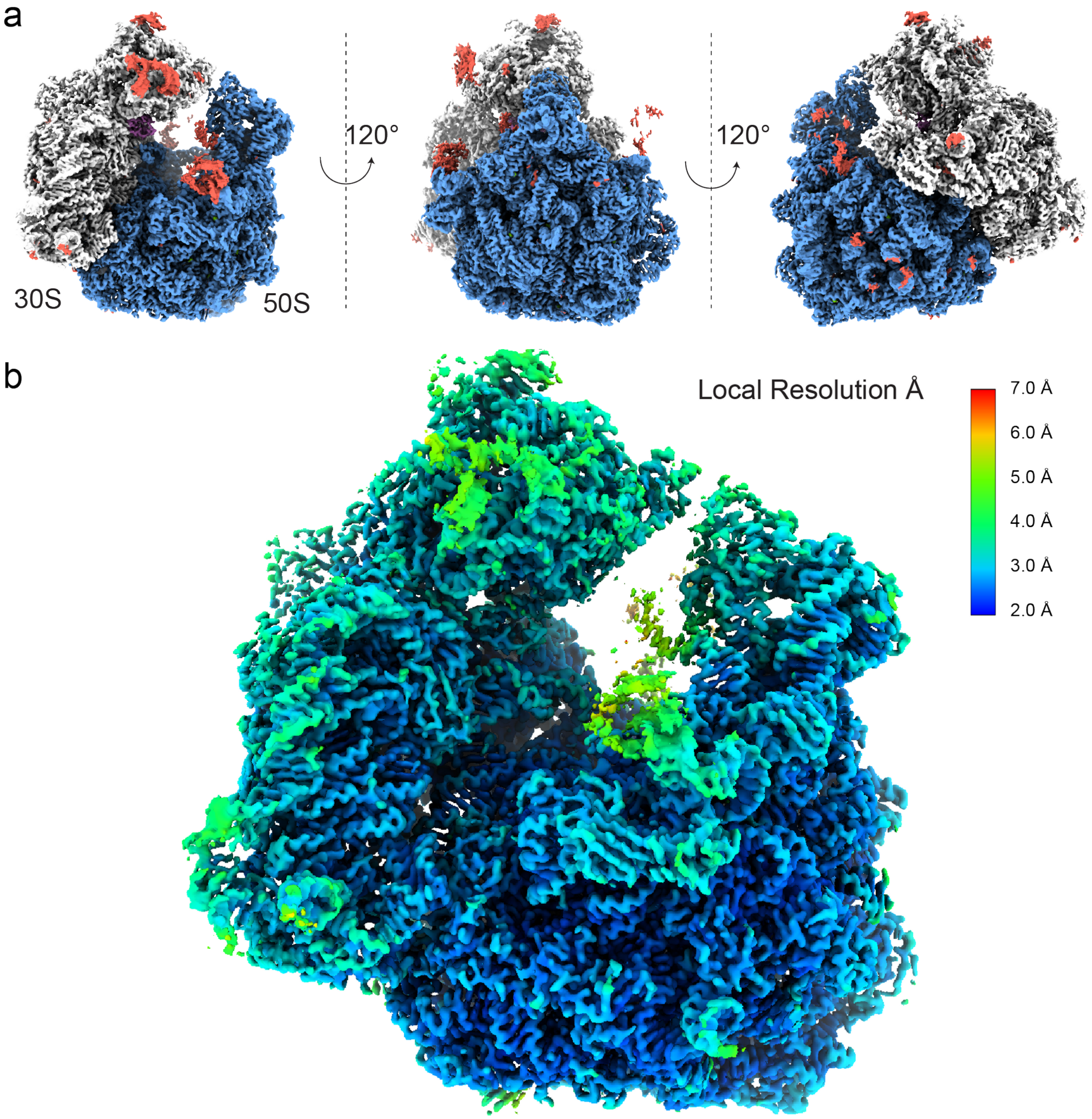

**SI Figure 9. CryoEM maps of 70S ribosome with Antibiotics Chloramphenicol and Gentamicin (Cm&GEN)**

**a** Cryo-EM maps coloured by chain, 30S white, 50S blue, RaiA violet. Maps with poor density that did not allow model building are shown in red. **b** Cryo-EM map coloured by local resolution estimation.

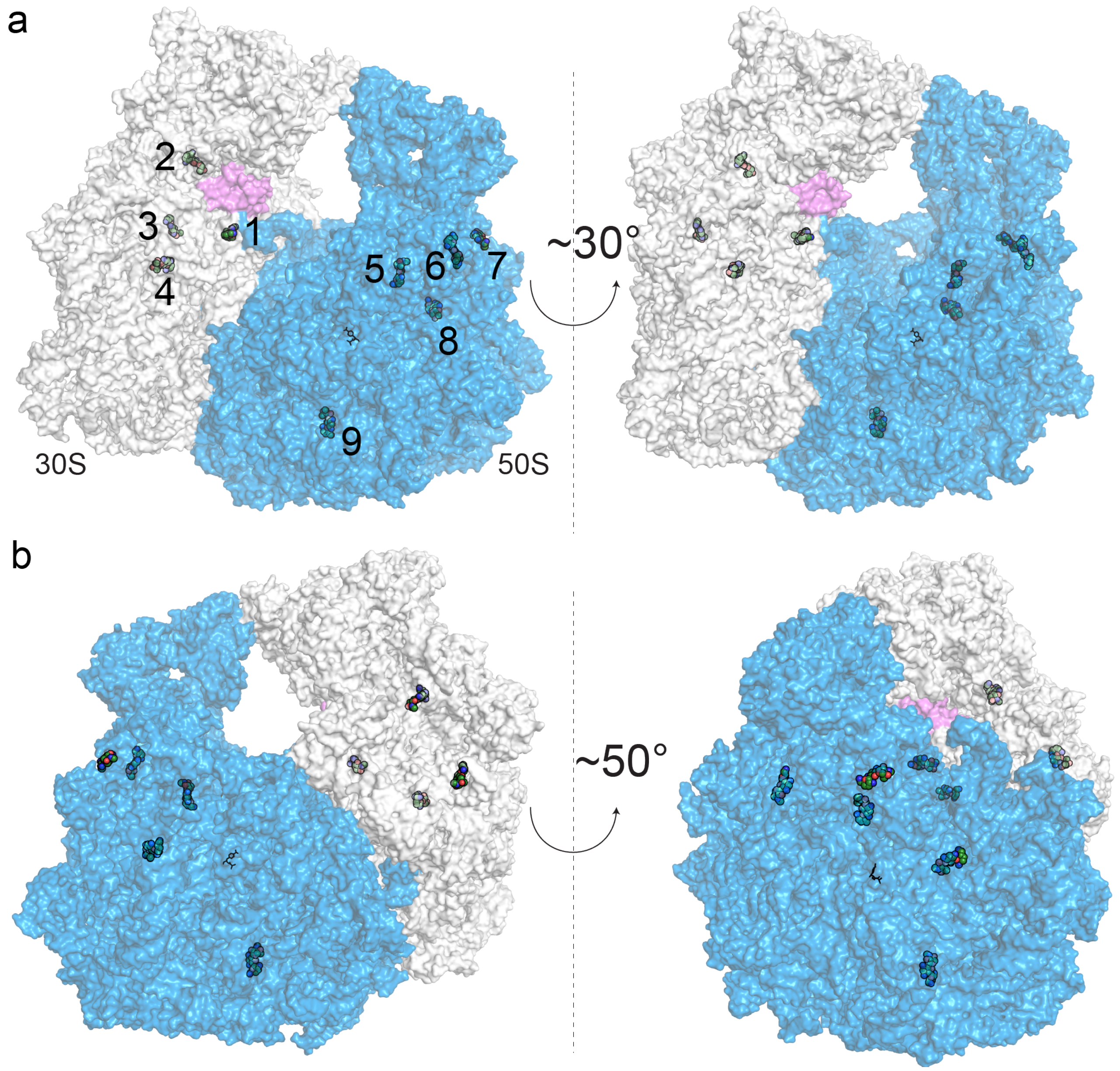

**SI Figure 10. CryoEM maps of 70S ribosome with Antibiotics**
**a** Cryo-EM maps coloured by chain, 30S white, 50S blue, RaiA violet. Maps with poor density that did not allow model building are shown in red. **b** Cryo-EM map coloured by local resolution estimation.

**Supplementary Table 1: Cryo-EM data collection, refinement and validation statistics.**

| | *F. tularensis* 70S (EMDB-54782) (PDB- **9SDA**) | *F. tularensis* 70S CM&GEN (EMDB-55615) (PDB- **9T6H**) |
|---|---|---|
| **Data collection and processing** | | |
| Microscope | Titan Krios | Titan Krios |
| Detector | Falcon 4i | Falcon 4i |
| Magnification (nominal) | 165.000x | 165.000x |
| Voltage (kV) | 300 | 300 |
| Spherical aberration | 2.7 mm | 2.7 mm |
| Total electron dose (e–/Å$^2$) | 40 | 60 |
| Defocus range (μm) | -3.0 to -0.8 | -3.0 to -0.8 |
| Pixel size (Å) | 0.76 | 0.76 |
| Stage tilt | 0° | 0° |
| Number of Micrographs | 28,005 | 5468 |
| Final particle images (no.) | 476,383 | 149,692 |
| Map resolution (Å) [FSC threshold] | 2.50 [$FSC_{0.143}$] | 2.55 [$FSC_{0.143}$] |
| **Refinement** | | |
| Initial model used (PDB code) | 9SDA | 9T6H |
| Symmetry during reconstruction | C1 | C1 |
| RMSD | | |
| Bond lengths (Å) | 0.007 | 0.007 |
| Bond angles (°) | 0.817 | 0.818 |
| Validation | | |
| MolProbity score | 1.10 | 1.12 |
| Clashscore, all-atom | 1.80 | 1.93 |
| Rotamer outliers | 0.55% | 0.55% |
| Ramachandran plot | | |

| | | |
|---|---|---|
| Favored | 97.07% | 97.07% |
| Allowed | 2.93% | 2..93% |
| Outliers | 0.00% | 0.00% |

| **Model vs. Data** | | |
|---|---|---|
| Ligands (no.) | 0 | 10 (1xCm, 9xGEN) |
| $CC_{unsharpened}$ (mask/box/ligand) | 0.93 / 0.86 / -- | 0.86 / 0.82/ 0.74 |
